# α-/γ-Taxilin are required for centriolar subdistal appendage assembly and microtubule organization

**DOI:** 10.1101/2021.09.08.459392

**Authors:** Dandan Ma, Rongyi Wang, Fulin Wang, Zhiquan Chen, Ning Huang, Yuqing Xia, Junlin Teng, Jianguo Chen

## Abstract

The centrosome, composed of a pair of centrioles (mother and daughter centrioles) and pericentriolar material, is mainly responsible for microtubule nucleation and anchorage in animal cells. The subdistal appendage (SDA) is a centriolar structure located at the subdistal region on the mother centriole, and it functions in microtubule anchorage. However, the molecular composition and detailed structure of SDA remain largely unknown. Here, we identified a-taxilin and r-taxilin as new SDA components, which form a complex via their coiled-coil domains and serve as a new subgroup during SDA hierarchical assembly. Their SDA localization is dependent on ODF2, and α-taxilin recruits CEP170 to the SDA. Functional analyses suggest that α-taxilin and γ-taxilin are responsible for centrosomal microtubule anchorage during interphase, as well as for proper spindle orientation during metaphase. Altogether, our results shed light on the molecular components and functional understanding of the SDA hierarchical assembly and microtubule organization.

## Introduction

The centrosome, the main microtubule organizing center (MTOC) in animal cells, participates in microtubule-related activities that include cell division (*Cabral et al., 2019*), cell polarity maintenance (*Burute et al., 2017*), cell signaling transduction (Barvitenko et al., 2008), and ciliogenesis (*Pitaval et al., 2017*; *Tu et al., 2018*). The centrosome is a non-membrane bound organelle composed of a pair of orthogonally arranged centrioles and surrounded by pericentriolar material (PCM) (*Delattre and Gonczy, 2004*). The two centrioles, mother and daughter centrioles, can be distinguished from each other by the decorations at the distal and subdistal ends of the mother centriole, called the distal/subdistal appendages (DAs/SDAs).

Accumulated data have revealed the structure characteristics and functions of DAs and SDAs. DA proteins, including CEP83, CEP89, FBF1, SCLT1, and CEP164, are essential for the centriole-to-membrane docking and TTBK2 recruitment that promote ciliogenesis (*Tanos et al., 2013*; *Huang et al., 2018*). Super-resolution microscopy has enabled researchers to identify the precise localization of the various centrosomal components, including the DA and SDA (*Yang et al., 2018*), and has revealed that SDAs differ from DAs in both molecular composition and intracellular functions (*Uzbekov and Alieva, 2018*). Located beneath the DAs, SDAs are composed of ODF2, CEP128, centriolin, NdelI, CCDC68, CCDC120, ninein, and CEP170 (*Huang et al., 2017*; *Kashihara et al., 2019*), which are classified into two groups according to their specific localizations: the ODF group (ODF2, CEP128 and centriolin), which is localized on SDAs, and the ninein group (ninein, Kif2a, p150glued, CCDC68, CCDC120 and CEP170), localized on both SDAs and the proximal ends of centrioles (*Mazo et al., 2016*; *Huang et al., 2017*). Of them, ODF2 occupies the bases of both the DAs and SDAs via different domains, thus participating in the hierarchical assembly of both DA and SDA (*Tateishi et al., 2013*; *Chong et al., 2020*). Also, ODF2 ensures that microtubules focus properly at the centrosome, which then controls microtubule organization and stability (*Hung et al., 2016a*), and centriolin, ninein, CCDC68 and CCDC120 are all required for microtubule stabilization and anchorage at centrosomes (*Gromley et al., 2003*; *Mogensen et al., 2000*; *Huang et al., 2017*). The SDA proteins cooperate to form a microtubule anchoring complex that maintains the microtubule array. Although this complex has been well described, most likely there are other unidentified SDA proteins still exist.

The taxilin family includes α-taxilin, β-taxilin and γ-taxilin, which were reported to interact with syntaxin family members and participate in intracellular vesicle transport (*Nogami et al., 2003*). Here, we have identified α- and γ-taxilin as new SDA structural components. They are recruited to SDAs via ODF2, form ring-like structures between CCDC120 and ninein, and participate in CEP170 assembly. Like their colleagues in the SDAs, α- and γ-taxilin are involved in microtubule anchorage at the centrosome and are also indispensable for spindle orientation during metaphase.

## Results

### Screening of α-taxilin and γ-taxilin as new SDA components

To search for new SDA components, we labeled two SDA components, CCDC68 and CCDC120, which we had identified in a previous study (*Huang et al., 2017*) by infusing them with V5-tagged APEX2, and established CCDC68 and CCDC120 proximal candidates via the mass spectrometry analysis of the APEX2-mediated biotinylation (*Hung et al., 2016b*).

Among those candidates for CCDC68 and CCDC120 proximal labeling, we found a series of previously identified centriolar proteins such as γ-tubulin and STIL, thus verifying the efficiency of our centrosomal proteomics approach (*Figure 1—figure supplement 1A*). Since several centrosomal proteins possess the coiled-coil domain that is the basis for the protein-protein interactions essential for centrosome conFigureuration (*Andersen et al., 2003*), we focused on proteins containing one or more coiled-coil domains. To determine their sub-cellular localizations, each protein candidate was tagged (e.g., V5, mNeonGreen, or pmEmerald) and expressed in U2OS cells. Other than the already known centrosome proteins, five new proteins including α-taxilin, DACT1, DRG2, NCAPH2, and SMAP2 (*Figure 1—figure supplement 1A-B*), were identified to be co-localized with centrosome markers CP110 or γ-tubulin, thus proving that they resided on centrosomes. Among them, DRG2 was found in both CCDC68 and CCDC120 proximal candidates; α-taxilin, NCAPH2, and SMAP2 were found in CCDC68 proximal candidates; and DACT1 was found in CCDC120 proximal candidates (*Figure 1—figure supplement 1A-B*). γ-Taxilin was previously reported to be co-localized with Nek2A at the centrosome during interphase (*Makiyama et al., 2018*), here we found it co-localized with γ-tubulin, and also as CCDC68 and CCDC120 proximal candidates (*Figure 1—figure supplement 1A-B*), confirming its centrosomal localization.

**Figure 1.**
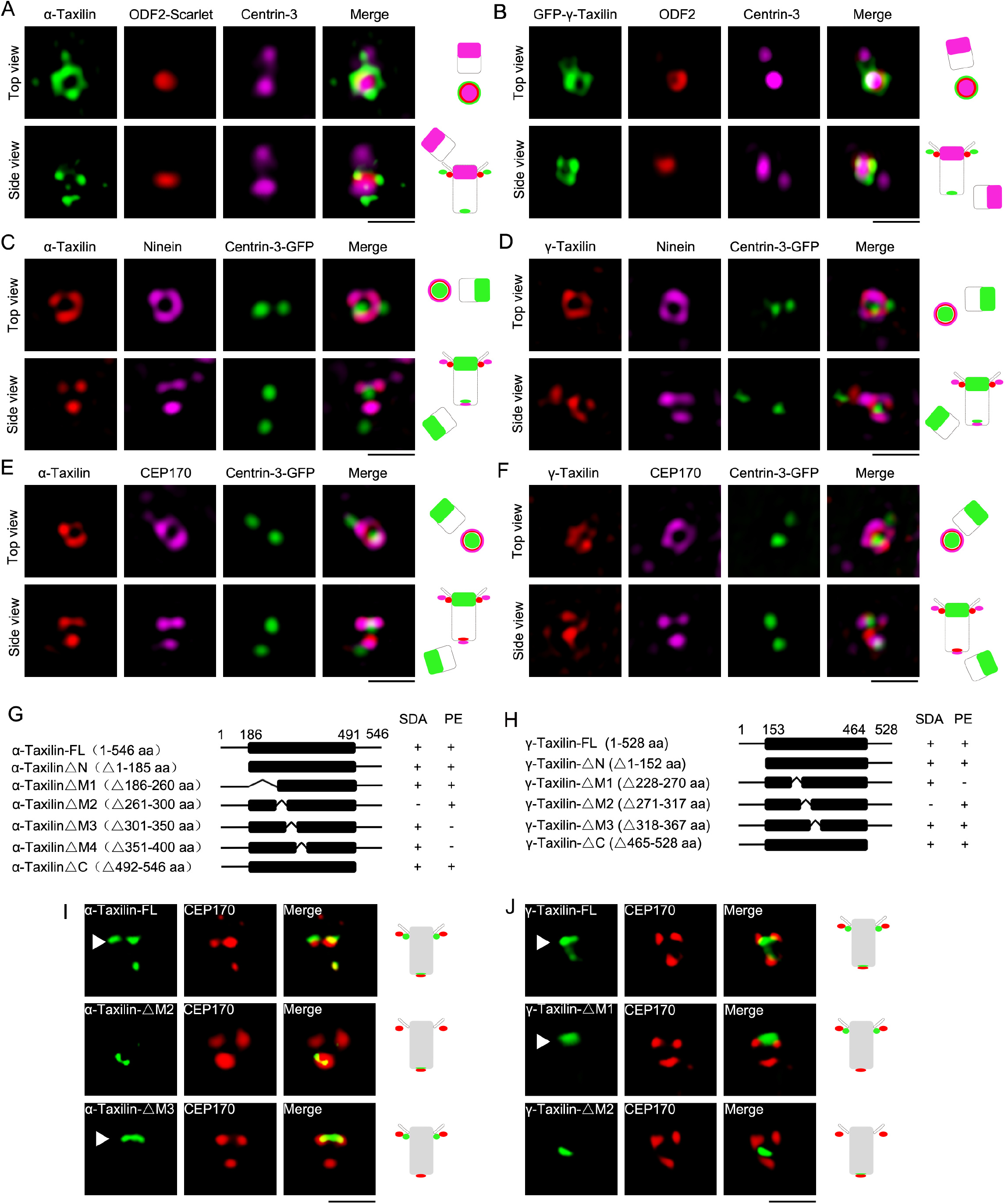
Structured illumination microscopy (SIM) images and characterizations of α-taxilin and γ-taxilin at the centrosomes. **(A)** Immunostained α-Taxilin (green) and centrin-3 (magenta) in RPE-1 cells transfected with ODF2-Scarlet (red). Scale bar, 1 µm. The cartoons to the right of each set of images graphically depict the merge images. **(B)** Immunostained ODF2 (red) and centrin-3 (magenta) in RPE-1 cells transfected with GFP-γ-taxilin (green). Scale bar, 1 µm. **(C)** Immunostained α-Taxilin (red) and ninein (magenta) in RPE-1 cells transfected with centrin-3-GFP (green). Scale bar, 1 µm. **(D)** Immunostained γ-Taxilin (red) and ninein (magenta) in RPE-1 cells transfected with centrin-3-GFP (green). Scale bar, 1 µm. **(E)** Immunostained α-Taxilin (red) and CEP170 (magenta) in RPE-1 cells transfected with centrin-3-GFP (green). Scale bar, 1 µm. **(F)** Immunostained γ-Taxilin (red) and CEP170 (magenta) in RPE-1 cells transfected with centrin-3-GFP (green). Scale bar, 1 µm. **(G-H)** Schematic showing full-length (FL) and deletion mutants (Δ) of α-taxilin (G) and γ-taxilin (H). N, N terminus; M, middle; C, C terminus; +, positive; −, negative; SDA, subdistal appendages; PE, proximal end. **(I)** Immunofluorescence images of HA-tagged FL α-taxilin and deletion mutants (green) in (G) and CEP170 (red) in RPE-1 cells. Scale bar, 1 µm. **(J)** Immunofluorescence images of CEP170 (red) in RPE-1 cells transfected with GFP-tagged FL γ-taxilin and deletion mutants (green) in (H). Scale bar, 1 µm. Arrowheads in (I) and (J) show SDA α-taxilin and γ-taxilin localizations, respectively. **Figure 1—figure supplement 1.** Screen of α-Taxilin and γ-taxilin as subdistal appendage proteins by CCDC68 and CCDC120 proximal labeling, and their localization characteristics.

Among the proteins that showed centrosomal localization, α- and γ-taxilin are similar in that each has a long coiled-coil domain in their middle regions (*Figure 1—figure supplement 1C*) (*Makiyama et al., 2018*). Both the anti-α-taxilin antibody (Proteintech: 22357-1-AP) and anti-γ-taxilin antibody (Proteintech: 27558-1-AP), which were designed to recognize the C-terminus of each of those proteins (*Figure 1—figure supplement 1C*), were chosen to detect their specificity. The results of western blotting showed that the molecular weights of these two proteins were about 72 kDa for α-Taxilin and 70 kDa for γ-taxilin as expected, respectively. The amount of those proteins decreased significantly after siRNA treatment (*Figure 1—figure supplement 1D*), indicating the antibodies’ specificity and the effectiveness of siRNAs.

Detailed α- and γ-taxilin localization at the centrosome was determined using super-resolution microscopy with a 3D-structured illumination system (SIM) (N-SIM, Nikon). In G1 phase, the top views of α- and γ-taxilin showed their ring-like patterns encompassing one centrin-3 dot, which resides in the distal lumen of centrioles (*Middendorp et al., 1997*) (*Figure. 1A-B*). The α- and γ-taxilin side views showed three dots occupying two levels with one level (two dots) beside ODF2 and thus at the SDA (*Figure. 1A-B*), and the other level (one dot) assumed to be at the proximal end (*Figure. 1A-E*). Those patterns resembled those of the ninein group, such as ninein and CEP170 (*Mazo et al., 2016*). Indeed, α-taxilin nearly co-localized with the ninein and CEP170 rings in the top view, and with the three ninein and/or CEP170 dots in the side view (*Figure. 1C, E*). Corresponding γ-taxilin patterns resembled those of α-taxilin by containing a smaller ring which resided inside the ninein and CEP170 signals (*Figure. 1D, F*).

We continued by asking which segments of α-taxilin and γ-taxilin were responsible for their SDA and proximal localizations. To examine that, we expressed a series of deletion mutants in RPE-1 cells and assayed their immunofluorescence using CEP170 as a marker (*Figure. 1G-H*). We found that the M2 deletion (261-300 aa) in α-taxilin eliminated its localization at the SDA, and deletion of M3 (301-350 aa) and M4 (351-400 aa) eliminated its localization at the proximal end (*Figure. 1I* and *Figure 1—figure supplement 1E*). Other deletion mutants (i.e., the N-terminus [1-185 aa], the C-terminus [492-546 aa] and M1 [186-260 aa]) did not affect its centrosomal localization pattern (*Figure 1-figure supplement 1E*). These data suggest that α-taxilin’s M2 (261-300 aa) is responsible for its SDA localization, while M3 (301-350 aa) and M4 (351-400 aa) work redundantly for its proximal end localization. For γ-taxilin, the M1 (228-270 aa) is responsible for its localization at the proximal end, while the M2 (271-317 aa) is responsible for its SDA localization (*Figure. 1J* and *Figure 1—figure supplement 1F*).

### The precise SDA localization of α-taxilin and γ-taxilin

Using stimulated emission depletion (STED) nanoscopy (TCS SP8 STED 3X, Leica), we further characterized α- and γ-taxilin rings with those of other SDA proteins and found that α- and γ-taxilin rings resembled those of other SDA proteins, including ODF2, CCDC68, CCDC120, ninein, and CEP170 (*Figure. 2A*). Of those, the ODF2 ring had the smallest diameter and the rest of those proteins’ rings increased in size in an order: CCDC68, CCDC120, γ-taxilin, α-taxilin, ninein, and CEP170 (*Figure. 2A-B* and *Table 1*). Therefore, those SDA proteins formed ordered concentric circles beginning with ODF2 and end with CEP170 following such an order: ODF2, CCDC68, CCDC120, γ-taxilin, α-taxilin, ninein and CEP170.

**Figure 2.**
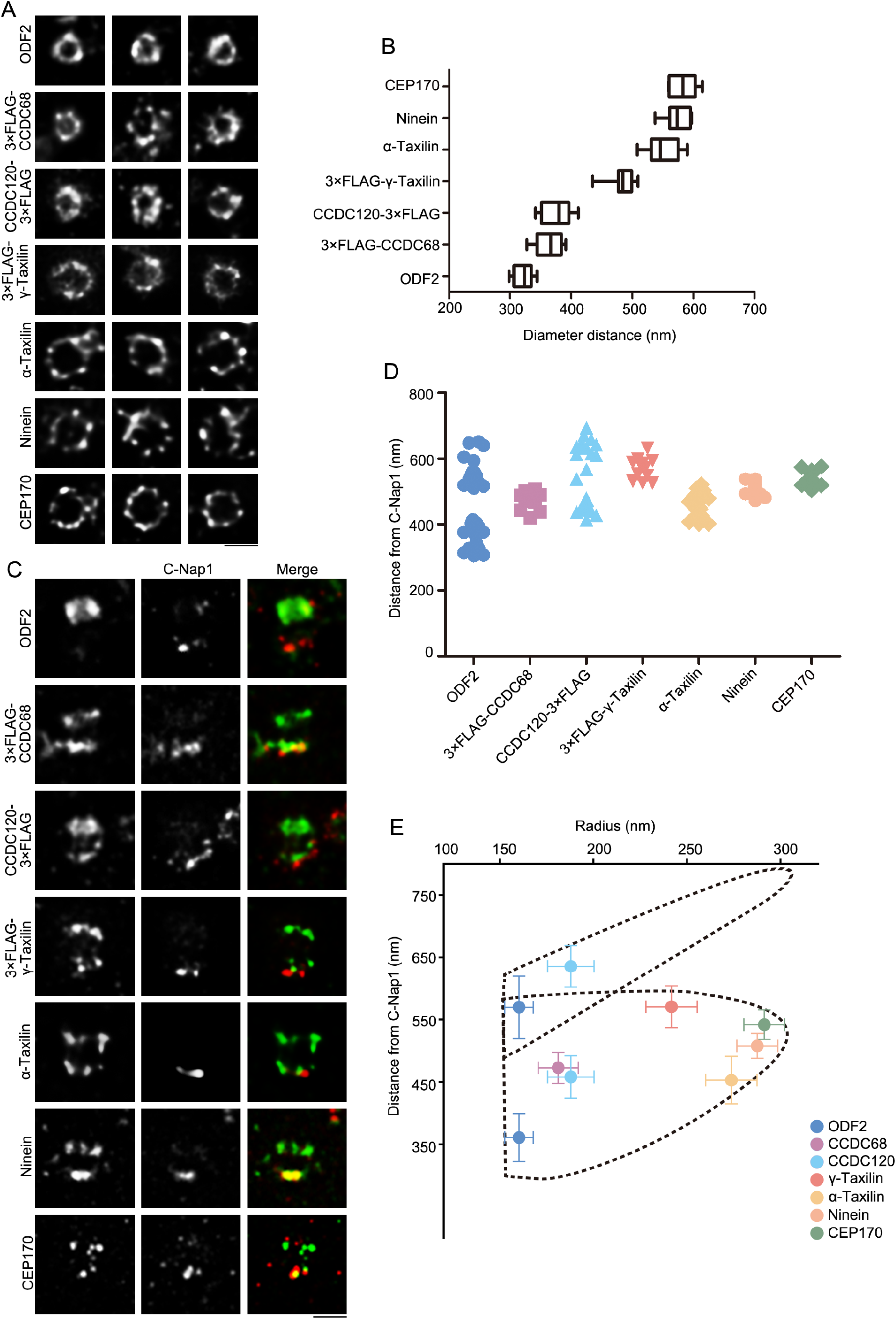
Super-resolved localizations of subdistal appendage (sDAP) proteins, including α-taxilin and γ-taxilin. **(A)** Representative simulated emission depletion (STED) super-resolution images showing top view of the subdistal appendage (SDA) protein distribution patterns. For ODF2, α-taxilin, ninein and CEP170, RPE-1 cells were immunostained with their antibodies. For CCDC68, CCDC120 or γ-taxilin, RPE-1 cells overexpressed with 3×Flag-tagged CCDC68, CCDC120 or γ-taxilin full-length were immunostained with Flag antibody. Scale bar, 500 nm. **(B)** Diameter analysis of SDA proteins showing ring size diameter. Data are mean (SD). n ≥ 7, box = 25^th^ and 75^th^ percentiles. (**C)** Representative two-color STED super-resolution images showing side view of the SDA proteins and the centriole proximal end protein C-Nap1. Scale bar, 500 nm. **(D)** A scatter plot describing the distance of sDAP proteins relative to C-Nap1. Data are mean (SD). n ≥11, box = 25^th^ and 75^th^ percentiles. **(E)** Relative localization of SDA proteins in radial and lateral directions of the mother centriole. The dotted lines revealing the slanted arrangement of distal appendage (DA) and the triangular structure of the SDA structure, respectively. Data are mean (SD).

**Table 1.** The diameter of subdistal appendage (SDA) proteins, including α-taxilin and γ-taxilin

| Protein | Diameter distance<br>(nm) | SD<br>(nm) | n |
| --- | --- | --- | --- |
| ODF2 | 320.59 | 15.26 | 14 |
| 3×FLAG-CCDC68 | 362.45 | 21.42 | 10 |
| CCDC120-3×FLAG | 375.90 | 19.64 | 9 |
| 3×FLAG-γ-Taxilin | 483.54 | 19.64 | 12 |
| α-taxilin | 547.47 | 27.44 | 7 |
| Ninein | 575.19 | 21.79 | 7 |
| CEP170 | 582.74 | 21.73 | 8 |

We then measured the longitudinal positions of SDA proteins by co-immunostaining with the proximal end protein, C-Nap1 (*Vlijm et al., 2018*), which serves as a position reference (*Figure. 2C-D*). ODF2 covered a wide range occupying between about 360 nm and 589 nm distances from C-Nap1, respectively, and was supposed to reside at the base of both DA and SDA (*Table 2*). This is comparable to the data obtained by the dSTORM super-resolution microscopy (*Chong et al., 2020*). Other SDA proteins all revealed two layered positions with one layer proximal with C-Nap1 and the other localizing between the two ODF2 layers, expect for CCDC120. The side view of CCDC120 showed three layered pattern, one at the proximal end, which resembled with the Ninein group proteins, while the other two population occupying about 458 nm and 636 nm distances from C-Nap1, and was assumed to reside at the SDA and DA structure, respectively (*Figure. 2C- D* and *Table 2*).

**Table 2.**
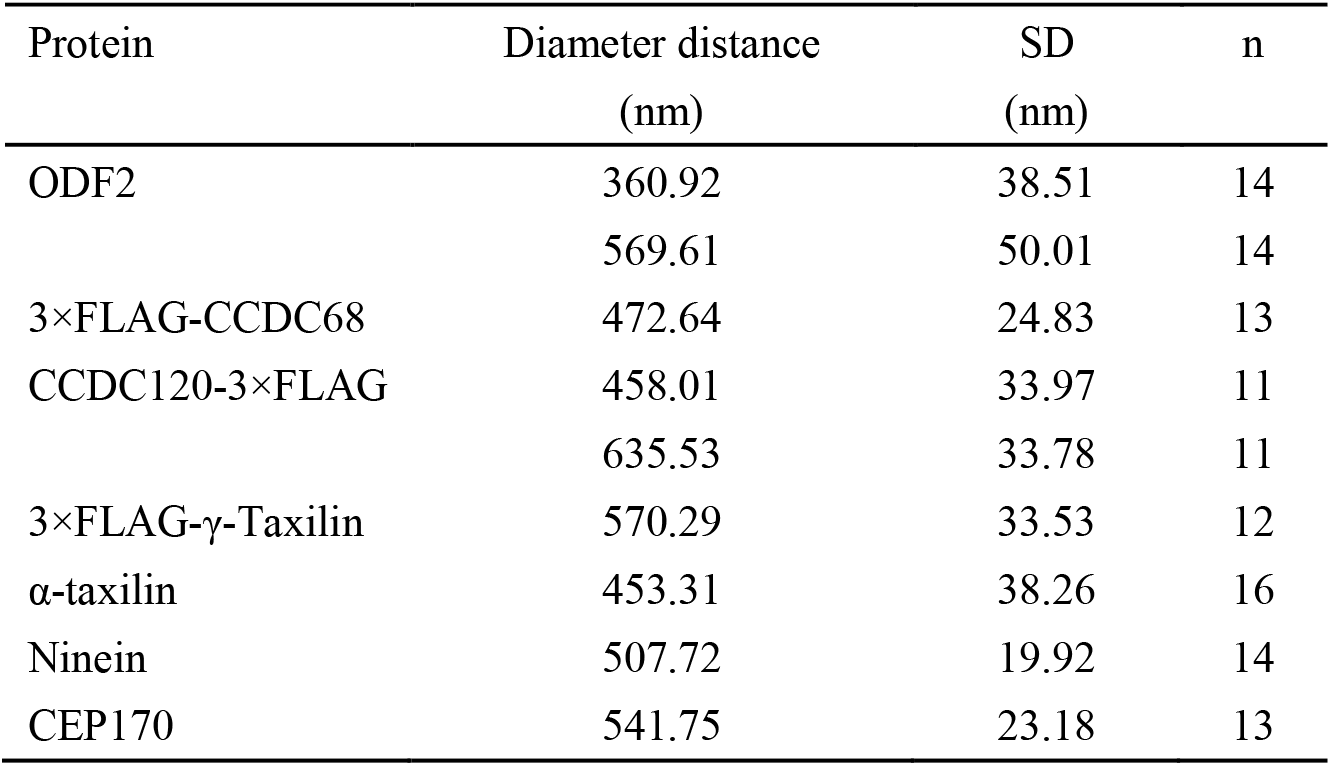
The longitudinal positions of subdistal appendage (SDA) proteins, including α-taxilin and γ-taxilin

Combining SDA measurements from the diameter and longitudinal positions from STED images, the relative localization of each SDA proteins was then established at the mother centriole. As was shown, ODF2 resides closest to the centriole wall, while ninein and CEP170 reside at the tip of SDA. At the SDA structure, CCDC68 and CCDC120 localize close to ODF2, and the α-taxilin and γ-taxilin located between CCDC120 and Ninein in concentric circles (*Figure. 2B, E*). In the longitudinal position, γ-taxilin was higher than CCDC68, CCDC120, Ninein and CEP170, while α-taxilin was lower than these proteins (*Figure. 2D-E*).

### α-Taxilin and γ-taxilin localization at the SDA depends on ODF2

To reveal the assembly manners of α- and γ-taxilin in the SDAs, the interactions between them and other SDA proteins were characterized through immunoprecipitation in human embryonic kidney (HEK)-239T and RPE-1 cells. First, ODF2 is located at the innermost layer of SDAs, and is the starting point for assembly of the other SDA components (e.g., TCHP, CCDC68, CCDC120, *etc*.) (*Ibi et al., 2011*; *Huang et al., 2017*). The results of immunoprecipitation revealed that endogenous ODF2 interacted with both α- and γ-taxilin in RPE-1 cells (*Figure. 3A-B*). Correspondingly, when we examined the specific localization of ODF2 and α- or γ-taxilin via STED nanoscopy, which is consistent with their specific localization as indicated in Figure. 2. We observe in the top view of the SDA region that α- and γ-taxilin encircled the ODF2 ring (*Figure. 3C-D*). When viewed from the side, we observed α- and γ-taxilin localized proximal to ODF2’s upper level (*Figure. 3C-D*).

**Figure 3.**
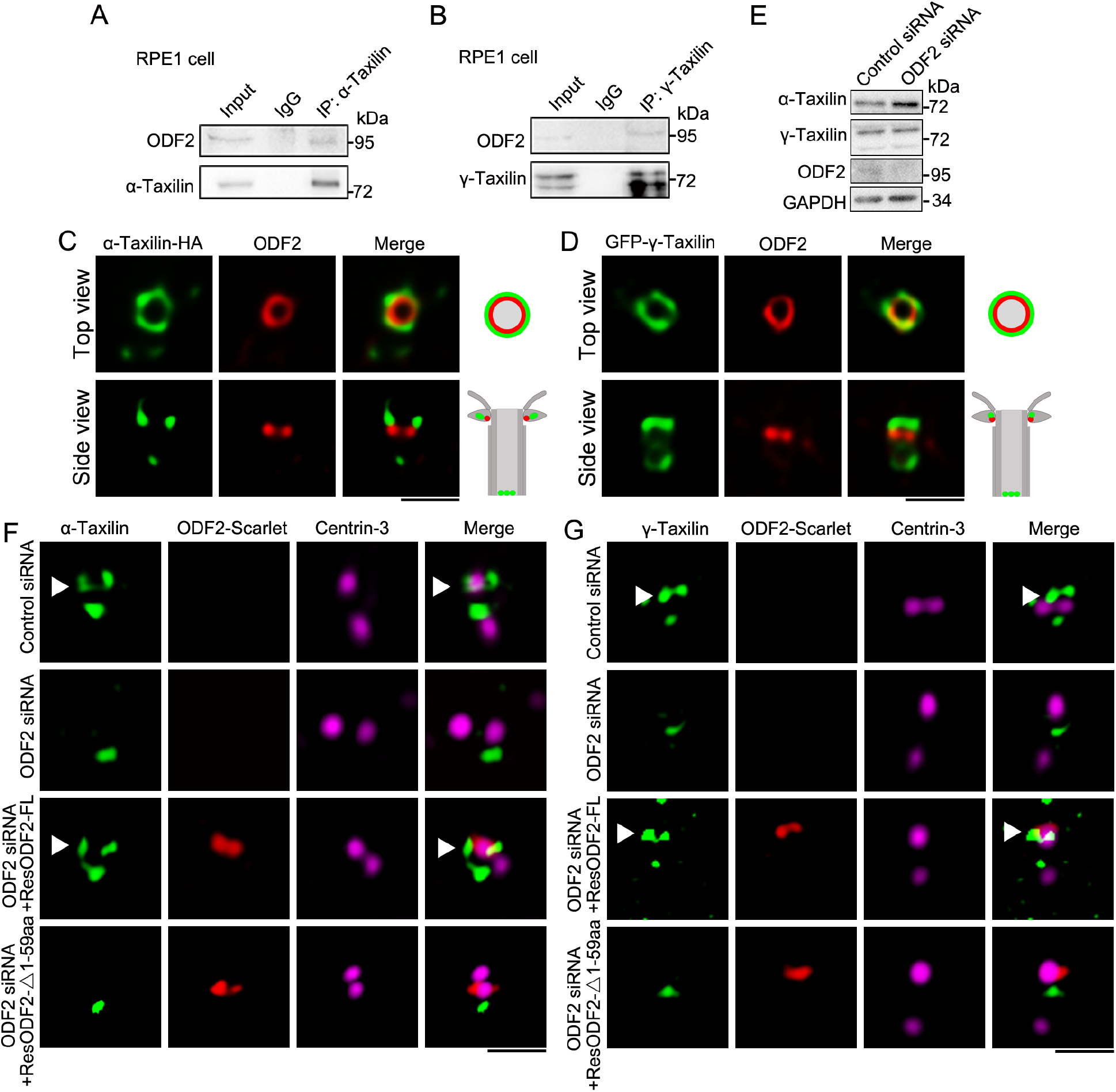
ODF2 is responsible for α-taxilin and γ-taxilin localization at the subdistal appendage (SDA). **(A)** Immunoblots of the endogenous immunoprecipitation (IP) assay of ODF2 and α-taxilin using anti-α-taxilin antibody in lysates of RPE-1 cells. IgG was the control. **(B)** Immunoblots of the endogenous IP assay of ODF2 and γ-taxilin using anti-γ-taxilin antibody in lysates of RPE-1 cells. **(C)** Simulated emission depletion (STED) images of RPE-1 cells immunostained with α-taxilin-HA (green) and ODF2 (red). Scale bar, 500 nm. **(D)** STED images of immunostained ODF2 (red) in RPE-1 cells transfected with GFP-**γ-**taxilin (green). The cartoons to the right of each set of images in (C) and (D) graphically depict the merge images. Scale bar, 500 nm. **(E)** Immunoblots of α-taxilin and γ-taxilin protein levels in control- or ODF2-siRNA treated RPE1 cells. GAPDH was the loading control. **(F)** 3D-structured illumination microscopy (SIM) images of immunostained α**-**taxilin (green) and centrin-3 (magenta) in control- or ODF2-siRNA treated RPE-1 cells and those cells rescued by transfection with either siRNA-resistant scarlet-tagged ODF2 full-length (FL) or 1-59 aa ODF2 deletion mutant (red). Scale bar, 1 µm. **(G)** 3D-SIM images of immunostained **γ-**taxilin (green) and centrin-3 (magenta) in control- or ODF2-siRNA treated RPE-1 cells and those cells rescued by transfection with either siRNA-resistant scarlet-tagged ODF2 FL or the 1-59 aa ODF2 deletion mutant (red). Scale bar, 1 µm. Arrowheads in (F) and (G) show α-taxilin and γ-taxilin SDA localizations, respectively. **Figure 3—figure supplement 1.** α-Taxilin and γ-taxilin assembly at the centrosome.

Furthermore, immunoblot analysis showed that when ODF2 was depleted by siRNA knockdown, α- and γ-taxilin levels in those cells remained unchanged (*Figure. 3E*). However, fluorescence intensity of both α- and γ-taxilin decreased at the centrosomes after ODF2 depletion (*Figure 3—figure supplement 1A-B*), suggesting that ODF2 affects their centrosomal localizations without affecting their protein levels. As shown by 3D-SIM images, ODF2 depletion by siRNA treatment resulted in α-taxilin or γ-taxilin signal loss at the SDA region, while their proximal end signals were not affected (*Figure. 3F-G*). Moreover, those lost SDA signals could be rescued by overexpressing full-length ODF2. However, the 1-59 aa deletion mutant, which affects ODF2 localization at SDA (*Tateishi et al., 2013*) and recruiment of other SDA components (*Huang et al., 2017*), could not rescue SDA localization of either α-taxilin or γ-taxilin after siRNA treatment (*Figure. 3F-G*). These data further illuminate the role of ODF2 in α- and γ-taxilin assembly at the SDAs.

In contrast, depleted CCDC68 or CCDC120 did not affect α-taxilin or γ-taxilin fluorescence intensity at centrosomes in RPE-1 cells (*Figure 3—figure supplement 1C-F*), and ectopically expressed CCDC68 and CCDC120 could not be immunoprecipitated from cell lysates by ectopically expressed α-taxilin or γ-taxilin in HEK-293T cells (*Figure 3—figure supplement 1G-J*). Previously, TCHP was reported to reside at the SDA midzone and to interact with ODF2 and ninein (*Ibi et al., 2011*). Here, endogenous TCHP did not interact with either α-taxilin or γ-taxilin in HEK-293T cells (*Figure 3—figure supplement 1K*). Therefore, the proper localization of α-taxilin and γ-taxilin at the SDA depends on ODF2, but not on CCDC68, CCDC120, or TCHP.

### α-Taxilin and γ-taxilin form a complex at SDA via their coiled-coil domains

We then moved on to determine the relationship between α-taxilin and γ-taxilin, which both belong to the taxilin family and each possesses a coiled-coil domain in the middle region (*Nogami et al., 2004*). First, using endogenous γ-taxilin and α-taxilin immunoprecipitation in HEK-293T cells, we detected an interaction between these two proteins (*Figure. 4A*). So, we then generated *α-taxilin* or *γ-taxilin* knockout (KO) RPE-1 cells using the CRISPR-Cas9 approach (*Figure 4****—****figure supplement 1A-B* and *Figure. 4B-C*). Compared with that of wild-type (WT) cells, α-taxilin fluorescence intensity was less at the *γ-taxilin* KO cell centrosomes (*Figure. 4D*). Conversely, *α-taxilin* depletion did not affect centrosomal γ-taxilin fluorescence intensity (*Figure. 4D*). Mapping with ectopically expressed γ-taxilin and α-taxilin truncated mutants indicated that they interact with each other via their coiled-coil domains (*Figure. 4E-H*). This was further confirmed via an *in vitro* binding assay using a purified bacteria-produced MBP-fused γ-taxilin M region (153-464 aa) and a GST tagged α-taxilin M region (186-491 aa) (*Figure. 4I*). These data suggest that γ-taxilin directly recruits α-taxilin to centrosomes via its coiled-coil domain.

**Figure 4.**
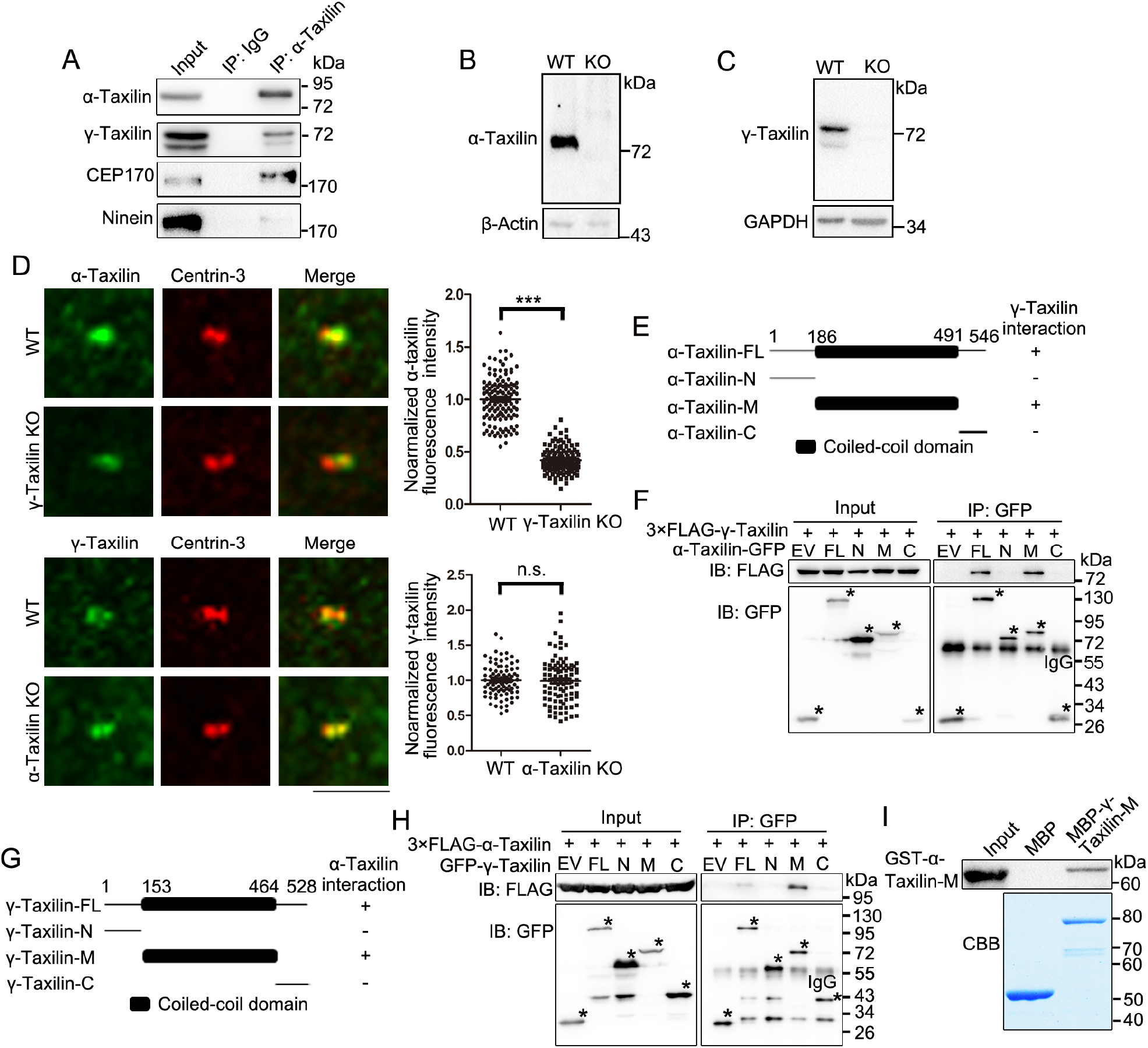
γ-Taxilin interacts with α-taxilin via their coiled-coil domains. **(A)** Endogenous immunoprecipitation (IP) assays of α-taxilin with γ-taxilin, ninein, and CEP170 using anti-α-taxilin antibody in lysates of HEK-293T cells and were immunoblotted with the indicated antibodies. IgG was the control. **(B-C)** Immunoblots showing depleted α-taxilin (B) and γ-taxilin (C) were in RPE-1 KO cells. β-Actin or GAPDH were used as loading controls. **(D)** Confocal images of immunostained α-taxilin (green) in wild-type (WT) and *γ-Taxilin* knockout (KO) RPE1 cells, as well as immunostained γ-Taxilin (green) in WT and *γ-Taxilin* KO RPE1 cells. Immunostained Centrin-3 (red) was used as a centrosome marker. Data are analyzed by two tailed Student’s *t*-test with three experimental replicates and expressed as mean (SEM). n > 60; ***, *p* < 0.001. n.s., not significant. **(E)** Schematic showing the α-taxilin full-length (FL) and truncated mutants’ (N terminus, Middle, and C terminus) interactions with γ-taxilin. +, positive; −, negative. **(F)** Lysates of HEK-293T cells co-overexpressing GFP-tagged FL α-taxilin or its truncated mutants from (E) with 3×FLAG-γ-Taxilin were subjected to IP and immunoblotted (IB) with anti-GFP and anti-FLAG antibodies. EV, empty vector. **(G)** Schematic showing FL γ-Taxilin and its truncated mutants’ interactions with α-taxilin. **(H)** Lysates of HEK-293T cells co-overexpressing GFP-tagged FL γ-taxilin or its truncated mutants from (G) with 3×FLAG-α-taxilin were subjected to IP with anti-GFP and IB with anti-GFP and anti-FLAG antibodies. **(I)** *In vitro* binding assay of MBP-γ-taxilin-M from (G) (expressed in *E. coli*, purified, and stained with Coomassie brilliant blue [CBB]) with GST-α-taxilin-M from (E) (expressed in *E. coli* and pulled down and detected by IB using the GST antibody). **Source data 1.** Data of centrosomal α-taxilin fluorescence intensity in wildtype (WT) and γ-taxilin knockout (KO) RPE1 cells. **Source data 2.** Data of centrosomal γ-Taxilin fluorescence intensity in wildtype (WT) and α-Taxilin knockout (KO) RPE1 cells. **Figure 4—figure supplement 1.** Partial sequences of *α-taxilin* or *γ-taxilin* knockout (KO) in RPE1 cells.

### α-Taxilin recruits CEP170 to SDAs

The SDA marker proteins ninein and CEP170 are located in the peripheral region of the SDA structure (*Huang et al., 2017*; *Chong et al., 2020*), and based on SDA protein localization pattern, α-taxilin located upstream of those two proteins (*Figure. 2A*). So, we investigated whether α-taxilin is involved in recruiting ninein and CEP170 to the SDAs. Endogenous immunoprecipitation assay showed that α-taxilin interacted with CEP170, but not with ninein (*Figure. 4A*). Also, α-taxilin and CEP170 co-localized at both the SDAs and the proximal ends, as observed with STED nanoscopy (*Figure. 5A*). To identify which segment of α-taxilin was responsible for that association, we overexpressed truncated mutants of HA-tagged α-taxilin in HEK-293T cells, and mapped the mutants’ interactions with CEP170. CEP170 interacted with α-taxilin in cells overexpressing full-length α-taxilin, as well as the M (186-491 aa) region and C-terminus (491-546 aa) deletion mutant constructs (ΔC), but not with the N-terminus construct (1-185 aa) (*Figure. 5B-C*). This suggests that the α-taxilin M region, but not the N- or C-terminal regions, is required for its interaction with CEP170. An *in vitro* binding assay using GST-tagged α-taxilin-M and 3×FLAG-CEP170 purified from bacteria and HEK-293T cells (*Figure. 5D*), respectively, suggested that α-taxilin may directly bind CEP170.

**Figure 5.**
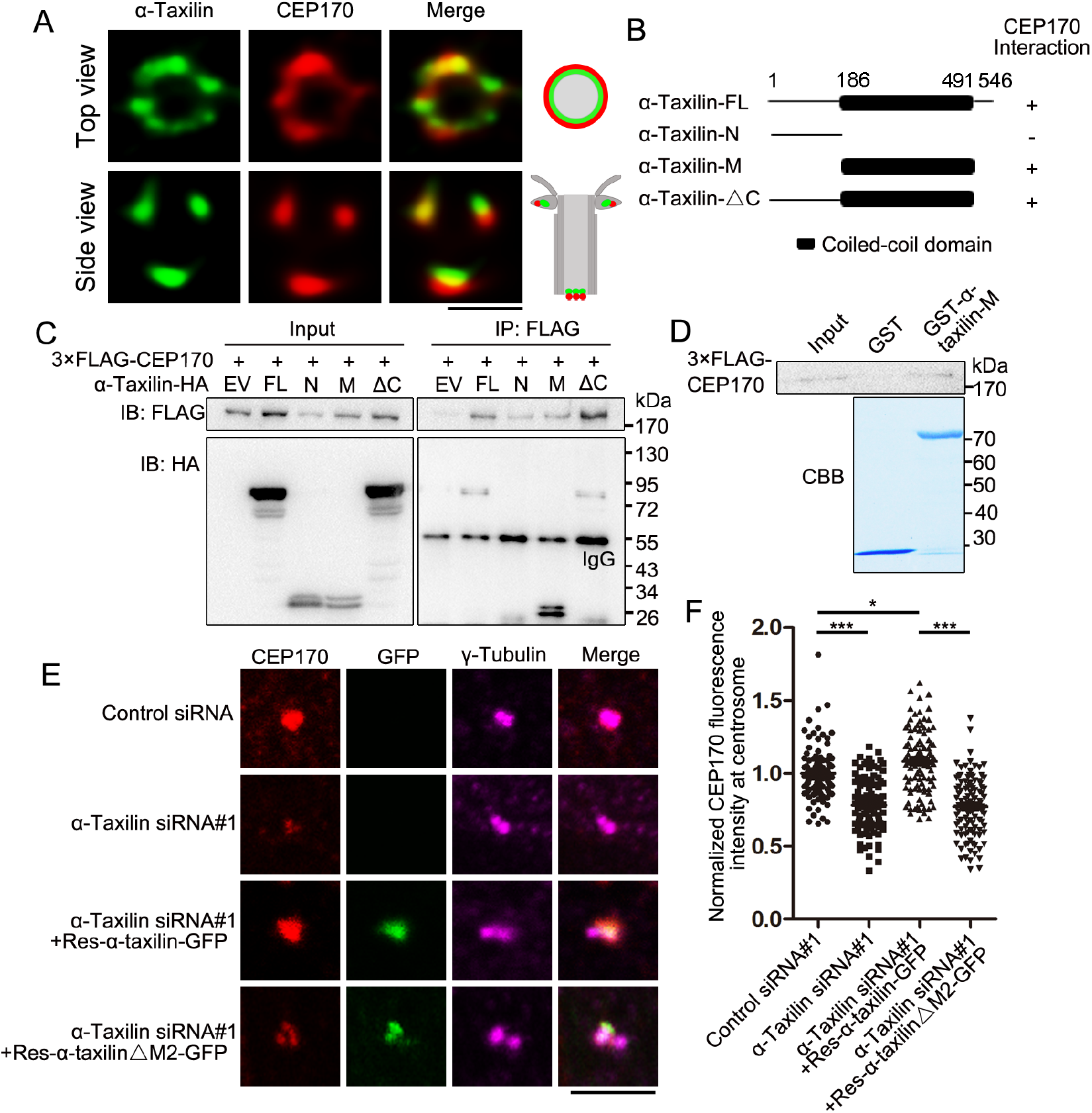
α-Taxilin recruits CEP170 to the subdistal appendage (SDA). **(A)** Simulated emission depletion (STED) images of RPE-1 cells immunostained with α**-**taxilin (green) and CEP170 (red). Scale bar, 500 nm. The cartoons to the right of the images graphically depict the merge images. **(B)** Schematic showing the full-length (FL) α-taxilin and the truncated mutants (N terminus, Middle, deleted C terminus). +, positive; −, negative. **(C)** Lysates of HEK-293T cells co-overexpressing HA-tagged α-taxilin-FL or the indicated truncated mutants in (B) with 3×FLAG-CEP170 were immunoprecipitated (IP) with anti-FLAG and immunoblotted (IB) with anti-HA and anti-FLAG antibodies. EV, empty vector. **(D)** *In vitro* binding of GST-α-taxilin-M from (B) (expressed in *E. coli* and purified) with 3×FLAG-CEP170 (expressed in HEK-293T cells and purified). The GST-α-taxilin-M was stained with Coomassie brilliant blue (CBB) and the 3×FLAG-CEP170 was pulled down and IB using FLAG antibody. **(E)** Confocal images of immunostained CEP170 (red) and γ-tubulin (magenta) in control- or α-taxilin-siRNA treated RPE-1 cells, and those cells rescued with siRNA-resistant GFP-tagged α-taxilin-FL and the α-taxilin M2 deletion mutant (Δ261-300 aa) (green). Scale bar, 5 µm. **(F)** Comparisons of CEP170 fluorescence intensities at the centrosomes in (E). Statistical significance was determined with one-way ANOVA with three replicates. Data are mean (SEM). n > 100; *, *p* < 0.05; ***, *p* < 0.001; n.s., not significant. **Source data 1.** Data of centrosomal CEP170 fluorescence intensity in control and α-taxilin siRNA treated RPE1 cells, and rescued by overexpression of α-Taxilin full-length or M2 deletion mutant.

Next, we examined the effect of α-taxilin depletion on CEP170’s centrosomal localization and detected a significant decrease in CEP170 fluorescence intensity in RPE-1 cells treated with α-taxilin siRNA, compared with control siRNA (*Figure. 5E-F*). Additionally, overexpressed siRNA-resistant full-length α-taxilin could rescue the CEP170 fluorescence intensity at the centrosomes, whereas the α-taxilin deletion mutant lacking the region responsible for its SDA localization (ΔM2) (Figure. 1G, I) failed (Figure. 5E-F). These indicate that α-taxilin participates with CEP170 in SDA assembly.

### α-Taxilin and γ-taxilin are responsible for centrosome microtubule anchoring during interphase

Since SDA serve mainly as microtubule anchoring sites at the centrosome, we examined whether α- and γ-taxilin are involved in microtubule organization. We began with a microtubule regrowth assay using *α-taxilin* or *γ-taxilin* KO RPE-1 cells (*Figure 4****—****figure supplement 1A-B* and *Figure. 4B-C*). After ice-induced microtubule depolymerization, an immunofluorescence assay detected microtubule dynamics after rewarming at 37 °C. RPE-1 cells were fixed at different periods of time (0, 5, and 10 min), during which microtubules formed arrays radiating from the centrosomes (*Figure. 6A-C*). The centrosomal microtubule aster in normal RPE-1 cells was visible 5 min after rewarming (*Figure. 6B*); and at 10 min, an extensive array of microtubules centering on the centrosome had formed (*Figure. 6C*). However, the microtubule asters were significantly impaired in the *α-taxilin* and *γ-taxilin* KO RPE-1 cells and the microtubule array densities were obviously less (*Figure. 6B-C*). To confirm this phenomenon, we further conducted microtubule regrowth assays in control and in α-taxilin and γ-taxilin siRNA-depleted RPE-1 cells and the results also displayed compromised microtubule reformation in the experimental cells, compared with that in the control cells (*Figure 6****—****figure supplement 1A-D*).

**Figure 6.**
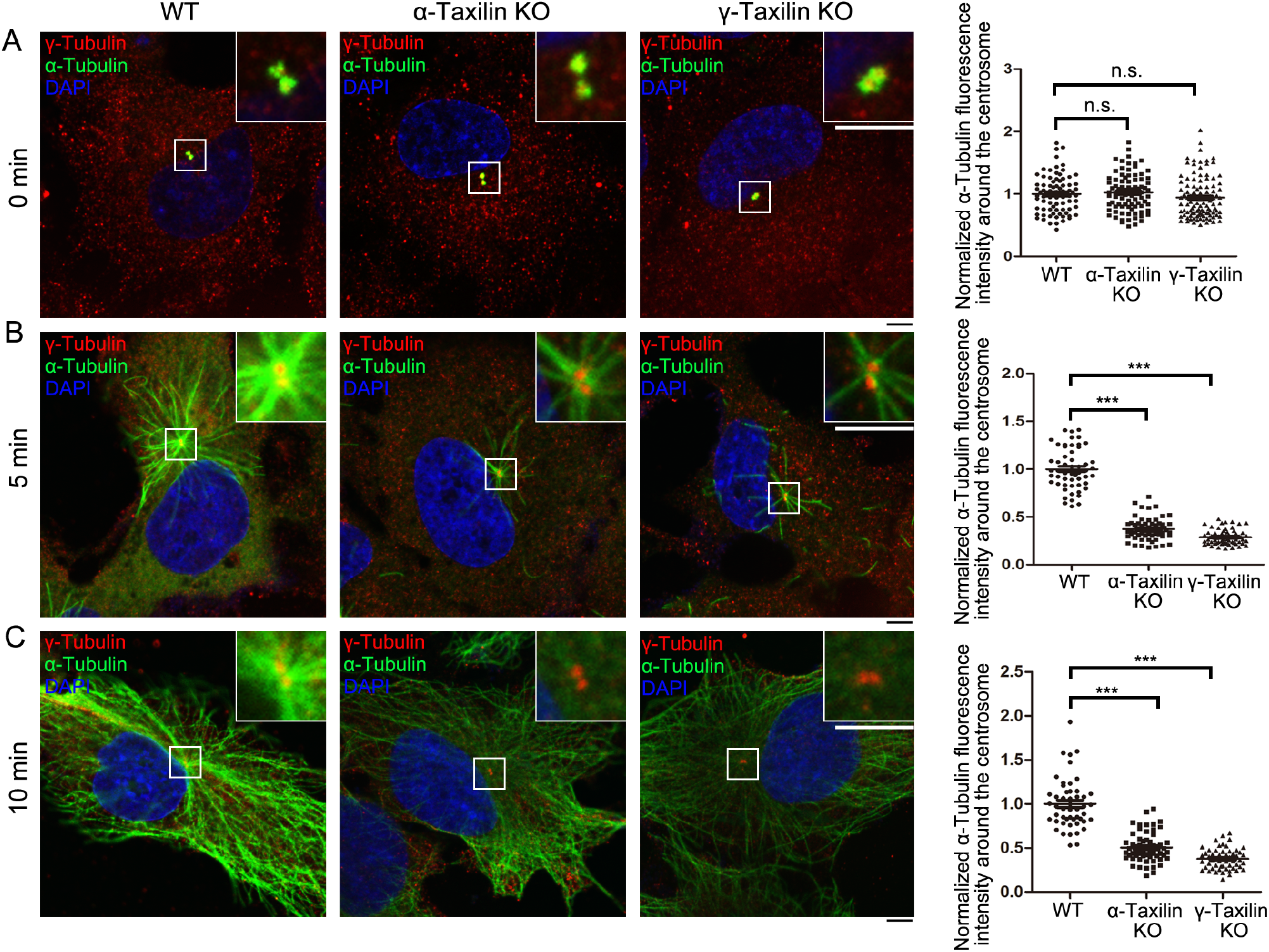
*α-Taxilin* and *γ-taxilin* knockout (KO) inhibits microtubule reformation after cold depolymerization. **(A-C)** Confocal images of microtubule reformation with immunostained α-tubulin (green) in wild-type (WT) and in *α-taxilin* or *γ-taxilin* KO RPE1 cells at 0 min (A), 5 min (B) and 10 min (C) after rewarming. Immunostained γ-tubulin (red) was the centrosome marker. DNA was stained with DAPI (blue). Scale bars, 5 µm. Statistical significance of α-tubulin fluorescence intensity at 0, 5, and 10 min was determined by one-way ANOVA. Data are mean (SEM). ***, *p* < 0.001; n.s., not significant. **Source data 1.** Data of normalized centrosomal α-tubulin fluorescence intensity in wildtype (WT), α-Taxilin and γ-taxilin knockout (KO) RPE1 cells. **Figure 6—figure supplement 1.** α-Taxilin and γ-taxilin depletion inhibits microtubule reformation in RPE-1 cells after cold depolymerization.

The centrosome acts as a MTOC by controlling microtubule nucleation and anchoring. γ-Tubulin, as a member of the γ-TuRC complex, plays a major role in microtubule nucleation (*Schatten and Sun, 2018*). To determine the causes of the compromised microtubule reformation observed in both α-taxilin and γ-taxilin depleted cells, we examined γ-tubulin at the interphase centrosome in control-, α-taxilin- or γ-taxilin-siRNA treated RPE-1 cells. Subsequently, we found that γ-tubulin fluorescence intensity at the interphase centrosome remained unchanged following both α-taxilin and γ-taxilin siRNA-induced depletion (*Figure 6****—****figure supplement 1E-F*), thus suggesting that microtubule reformation defects in α-taxilin or γ-taxilin depleted cells was caused by centrosome deficiency in microtubule anchoring instead of microtubule nucleation.

### α-Taxilin and γ-taxilin are required for proper spindle orientation during metaphase

In proliferating cells, centrosomes act as spindle poles when cells enter mitosis. Therefore, we investigated whether α-taxilin or γ-taxilin loss affected spindle formation. For this, we generated *α-taxilin* or *γ-taxilin* KO HeLa cells using the CRISPR-Cas9 approach (*Figure 7****—****figure supplement 1A-B* and *Figure. 7A-B*). Although bipolar spindle formation was not affected in *α-taxilin* or *γ-taxilin* KO HeLa cells (*Figure. 7C*), changes in the orientation of the spindle to the substratum were observed (*Figure. 7C-E*). Over half (51.7%) the *α-taxilin* KO cells showed greater than 10-degree spindle angles, while most spindle angles in WT cells were less than 5 degrees (55.9%). In the *γ-taxilin* KO HeLa cells, half (50.0%) of the cells had greater than 10-degree spindle angles, but similar spindle angles were found in only 8.3% of the WT cells (*Figure. 7D-E*). Spindle misorientations were rescued by overexpression of the 3×FLAG-tagged full-length α-taxilin or γ-taxilin (*Figure. 7C-E*), thus confirming their roles in maintaining proper spindle orientation during metaphase.

**Figure 7.**
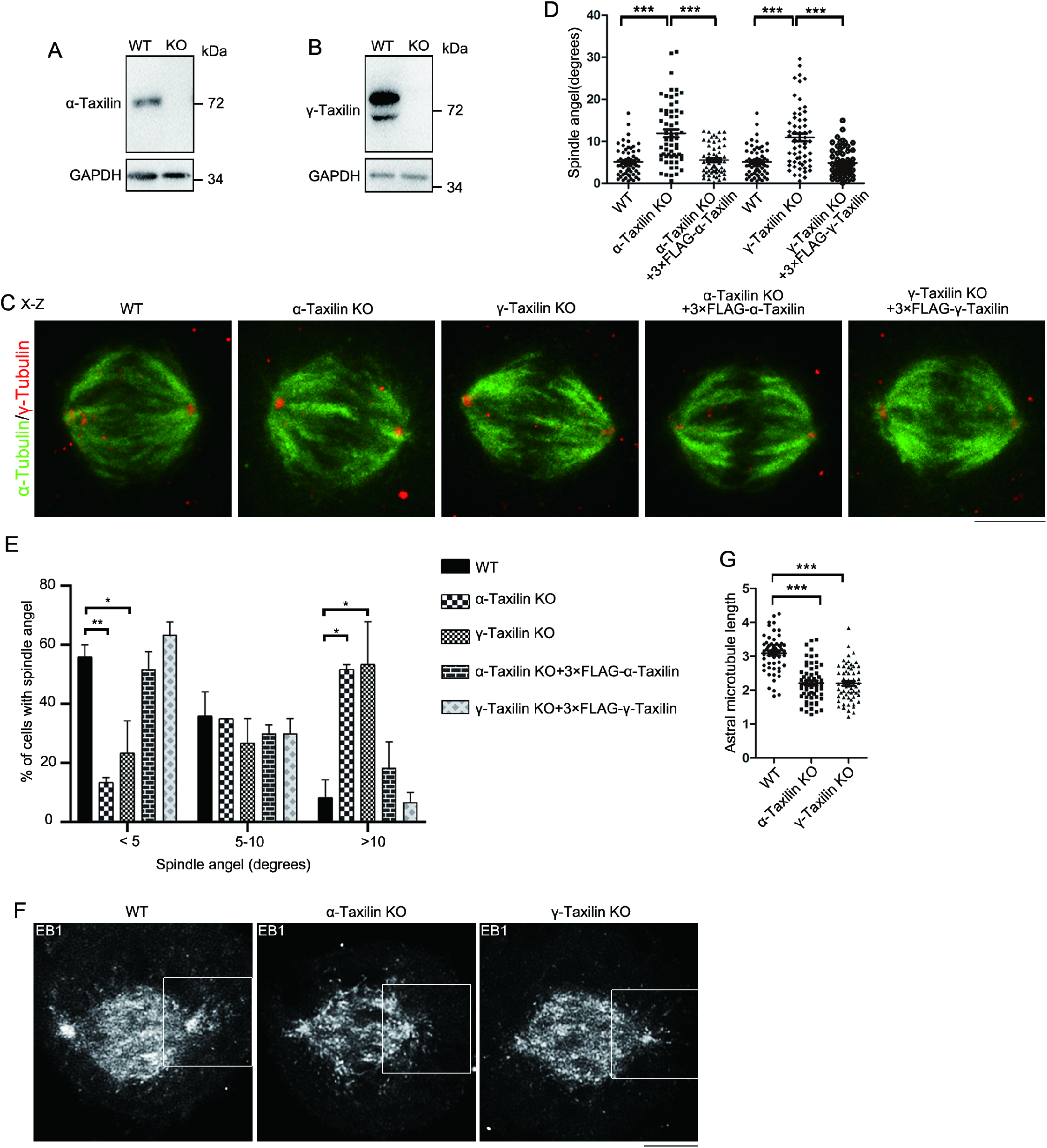
*α-Taxilin* and *γ-taxilin* knockout (KO) increases spindle angel orientation during mitosis. **(A-B)** Immunoblots showing depleted α-taxilin (A) and γ-taxilin (B) in respective KO HeLa cells. GAPDH was the loading control. **(C)** Representative orthogonal views (x-z) of metaphase HeLa cells stained for microtubules (α-tubulin, green) and spindle poles (γ-tubulin, red) in wild-type (WT) and in *α-taxilin* or *γ-taxilin* KO cells. Misoriented spindles were subsequently rescued by overexpression of 3×FLAG-tagged α-taxilin or γ-taxilin, respectively. Scale bars, 10 µm. **(D)** Comparisons of raw spindle angles (degrees) in WT, *α-taxilin* or *γ-taxilin* KO HeLa cells, and of those subsequently rescued with 3×FLAG-tagged α-taxilin or γ-taxilin. Data are represented as mean (SEM). Statistical significance was determined by one-way ANOVA of three individual experiments. n > 60. ***, *p* < 0.001. **(E)** Quantification of cells in (D) and comparisons of three spindle angle categories. Data are mean (SEM). Statistical significance was determined by one-way ANOVA of three independent experiments. n > 60. *, *p* < 0.05; **, *p* < 0.01. **(F)** Confocal images of aster microtubules immunostained with EB1 in WT, *α-taxilin* or *γ-taxilin* KO HeLa cells. Scale bars, 10 µm. **(G)** Comparisons of aster microtubule lengths in (F). Data are mean (SEM). Statistical significance was determined by one-way ANOVA of three replicates. n > 60. ***, *p* < 0.001. **Source data 1.** Data of spindle angels in wildtype (WT), α-Taxilin and γ-taxilin knockout (KO) HeLa cells, and rescued by overexpression of indicated full-length. **Source data 2.** Data of astral microtubule length in wildtype (WT), α-Taxilin and γ-taxilin knockout (KO) HeLa cells, and rescued by overexpression of indicated full-length. **Figure 7—figure supplement 1.** Partial sequences of *α-taxilin* or *γ-taxilin* knockout (KO) in HeLa cells.

Finally, because astral microtubule loss causes spindle misorientation (*Hung et al., 2016a*), we examined possible losses by staining the plus-end microtubule binding protein EB1 (*Honnappa et al., 2009*) in WT and in *α-taxilin* and *γ-taxilin* KO HeLa cells. While the WT aster microtubule length was 3.09 (SD 0.53) μm, those lengths in the *α-taxilin* (2.20 [0.54] μm) and *γ-taxilin* (2.19 [0.53] μm) KO HeLa cells were significantly less (*Figure. 7F-G*). These data suggest that the changes in spindle orientation observed in *α-taxilin* or *γ-taxilin* KO HeLa cells are likely caused by decreased astral microtubule length.

## Discussion

SDAs are conserved structures located at the subdistal end of the mother centriole, and their formation, together with that of the DAs, marks centriole maturity (*Uzbekov and Alieva, 2018*). They play important roles in microtubule organization and spindle arrangement, and participate in various biological processes such as cell division and cell differentiation (*Hall and Hehnly, 2021*). Here, we applied APEX2-mediated proximity-labeling to centrosome proteomics by using CCDC68 and CCDC120 as baits (*Hung et al., 2016b*; *Huang et al., 2017*). The results showed a variety of possible proteins, some of which are known centrosomal proteins like γ-tubulin and γ-taxilin, and several new centrosome-localized proteins were also identified (*Figure 1****—****figure supplement 1A-B*), such as α-taxilin, DACT1, NCAPH2, DRG2 and SMAP2. Among these proteins, α-taxilin and γ-taxilin are both located at the SDA and the proximal end of mother centriole (*Figure. 1-3*).

The SDA is a comprehensive structure that consists of a spreading radial distribution of concentric proteins, and SDA assembly follows an hierarchical relationship based on protein distributions (*Chong et al., 2020*). Comparing SDA protein diameters, those of α- and γ-taxilin were larger than those of CCDC68 and CCDC120, but smaller than those of ninein and CEP170. In the longitudinal position, γ-taxilin was relatively higher while α-taxilin was relatively lower than these SDA proteins. Associated with their relative localization with other SDA proteins, they were supposed to reside in the middle SDA zone (*Figure. 8A-B*).

**Figure 8.**
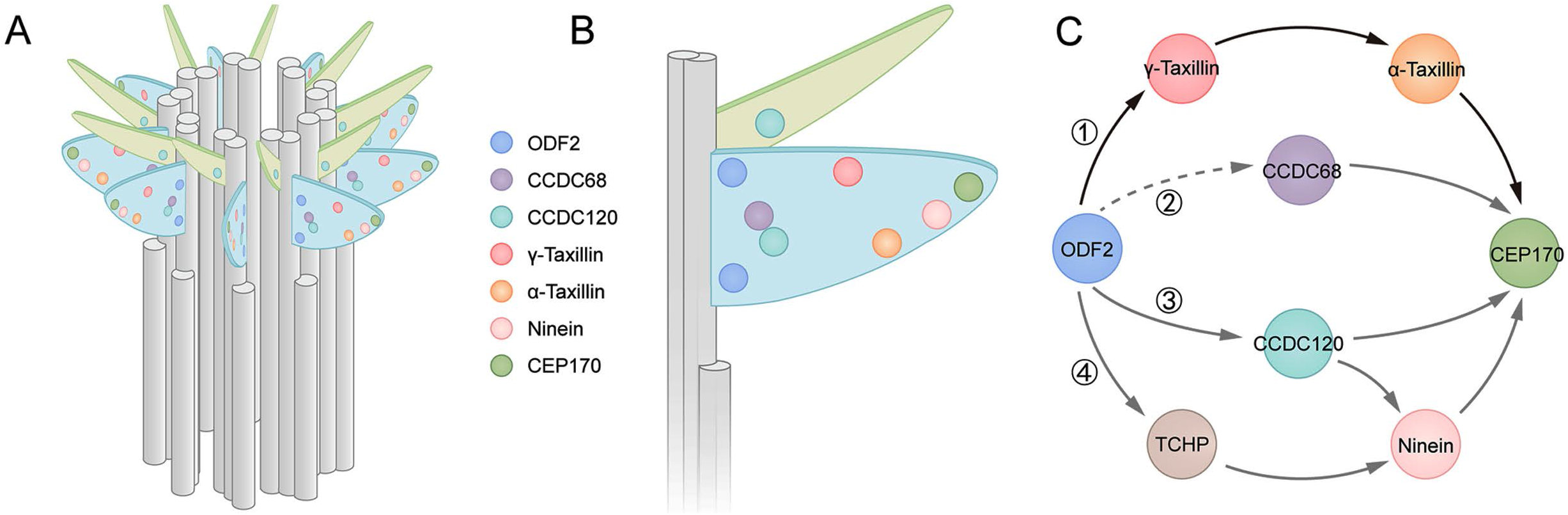
Models showing α-taxilin and γ-taxilin localization and assembly in the subdistal appendage (SDA). **(A)** A 3D model of a mother centriole, illustrating the localization of various SDA proteins, including ODF2, CCDC68, CCDC120, γ-taxilin, α-taxilin, ninein, and CEP170 in increasing diameter size order. **(B)** Close view of the SDA structure from (A). **(C)** Model showing the hierarchical relationships of SDA proteins with ODF2 at the SDA root. (1) ODF2 recruits γ-taxilin to the SDA, which in turn hierarchically recruits α-taxilin and then CEP170. (2) CCDC68 is recruited to the SDA, which further recruits CEP170 (*Huang et al., 2017*). (3) ODF2 recruits ninein and CEP170 via CCDC120 (*Huang et al., 2017*). (4) ODF2 recruits ninein via TCHP (Ihi et al., 2011). DA, distal appendages; SDA, subdistal appendages.

Among the currently known SDA components, ODF2 is the most closely located to the centriolar microtubules (*Figure. 8C*) and it likely recruits TCHP to assist ninein assembly at the SDA (*Ibi et al., 2011*). The ODF2 1-60 aa in its N-terminus is responsible for its association with CCDC120 (*Huang et al., 2017*). However, this association is not direct, as a yeast two-hybrid assay demonstrated. Besides, CCDC68 lies between ODF2 and CEP170, and how CCDC68 is recruited to the SDA is yet unknown (*Huang et al., 2017*). Our data suggest that ODF2 recruits α- and γ-taxilin, similarly to how it recruits CCDC120 (*Huang et al., 2017*). Since the ODF2 ring is much smaller than those of both γ-taxilin and α-taxilin, most likely the interactions between ODF2 and the taxilins are indirect (*Figure. 8A-B*). Although TCHP, CCDC68, and CCDC120 lie between ODF2 and the taxilins (*Figure. 8A-B*), no interactions among taxilins with those SDA proteins, except for ODF2, were detected by immunoprecipitation assays. Therefore, ODF2 recruits α-taxilin and γ-taxilin through a new pathway (*Figure. 8C*).

The functions of SDA components are critical for understanding this structure. Ninein is responsible not only for microtubule nucleation by docking the γ-TURC at the centrosomes (*Delgehyr et al., 2005*), but also for forming a microtubule-anchoring complex with CEP170 at the SDA periphery (*Pizon et al., 2020*). CCDC68 and CCDC120 also participate in microtubule anchoring (*Huang et al., 2017*). Similarly, compromised microtubule reformation following ice-depolymerization was observed in both α-taxilin and γ-taxilin depleted cells, likely because of depressed microtubule anchoring ability, as γ-tubulin intensity at the centrosome was not influenced by either α-taxilin or γ-taxilin siRNA knockdown. Therefore, our data suggest that α-taxilin and γ-taxilin serve as microtubule anchoring regulators at the centrosomes, functionally via a new pathway that is independent of CCDC68, CCDC120, and ninein.

In addition to acting as microtubule anchoring centers during interphase, SDAs also function as spindle regulators during mitosis. CEP170 interacts with centrosome-associated kinesins such as KIF2A, KIF2C, KIFC3, and spindle microtubule-associated KIF2B (Welburn et al., 2012; *Maliga et al., 2013*), which regulates spindle assembly and cell morphology. In our study, both α-taxilin and γ-taxilin depletion resulted in decreased astral microtubule length and increased spindle misorientation. As the hierarchical assembly of γ-taxilin, α-taxilin, and CEP170 was established, we speculate that α-taxilin and γ-taxilin regulate spindle orientation mainly by controlling CEP170’s centrosomal localization. In addition to CEP170, the most upstream SDA protein, ODF2, was shown to regulate spindle orientation via microtubule organization and stability (*Hung et al., 2016a*). Spindle misorganization has been extensively correlated with cell differentiation, cancer and neurological diseases such as microcephaly and lissencephaly (*Noatynska et al., 2012*), as well as tubular organ diseases (*Zhong and Zhou, 2017*). Whether α- and γ-taxilin are involved in development and diseases needs further investigation.

In conclusion, our results show α-taxilin and γ-taxilin to be new SDA proteins and shed new light on how SDAs are assembled. However, other yet unknown components exist at the SDA upper zone and participate in SDA assembly, and they and their functions need to be identified. Additionally, how α-taxilin and γ-taxilin function in development and diseases warrants further investigation.

## Materials and Methods

### Cell cultures and transfection

HeLa (ATCC, CCL-2), U2OS (ATCC, HTB-96), and HEK-293T (ATCC, CRL-3216) cells were purchased from ATCC (Manassas, VA, USA) and cultured in DMEM (GIBCO, Thermo Fisher Scientific, Waltham, MA, USA) with 10% FBS (CellMax, Lanzhou Minhai Bio-Engineering, Lanzhou, China) and RPE-1 cells (ATCC, CRL-4000) in DMEM/F12 (1:1) (GIBCO) with 10% fetal bovine serum, then all cells were cultured at 37 °C with a 5% CO_2_ atmosphere. The HeLa, U2OS, and HEK-293T cells were transfected using polyethylenimine (#23966-1, Polysciences), while the RPE-1 cells were transfected using Lipofectamine 3000 (Thermo Fisher Scientific) according to the manufacturer’s instructions.

### Plasmid construction

To obtain CCDC68-V5-APEX2 and CCDC120-V5-APEX2 constructs, human CCDC68 (NM_025214.3) and CCDC120 (NM_001163321.4), respectively, were amplified by PCR from human cDNA and cloned into mito-V5-APEX2 (Lam et al., 2015). Mito-V5-APEX2 was a gift from Alice Ting (Addgene plasmid # 72480; http://n2t.net/addgene:72480; RRID: Addgene_72480). Human γ-taxilin (XM_024452398.1), SWAP2 (AY358944.1), and DACT1 (NM_016651.6) were amplified by PCR from human cDNA and cloned into pmNeonGreenHO-G (*Tanida-Miyake et al., 2018*). pmNeonGreenHO-G was a gift from Isei Tanida (Addgene plasmid # 127912; http://n2t.net/addgene:127912; RRID:Addgene_127912). Human ODF2 (NM_001242352.1) and its deletion mutant (Δ1-59 aa) were amplified by PCR from human cDNA and cloned into pScarletN1. pScarlet-N1 was a gift from Oskar Laur (Addgene plasmid # 128060; http://n2t.net/addgene:128060; RRID:Addgene_128060). Using PCR, α-taxilin (NM_175852.4) and its truncated segments were amplified from human cDNA and cloned into pcDNA3.1(+) (Sigma-Aldrich). Full-length α-taxilin and γ-taxilin were also cloned into p3×FLAG-CMV-14 (Sigma-Aldrich). Ninein-GFP, 3×FLAG-CEP170, and CCDC120-GFP were obtained from a previous study (*Huang et al., 2017*).

### APEX2-mediated proximal labeling of CCDC68 and CCDC120

Once CCDC68-V5-APEX2 and CCDC120-V5-APEX2 were constructed, a large-scale proteomic experiment was conducted in cultured HEK-293T cells in which both constructs were over-expressed for 24 h. Following treatment with BP for 30 min and H_2_O_2_ for 1 min, the biotinylated proteins were enriched by Streptavidin-coated beads for mass spectrographic analysis and the resulting raw data were then searched in the Uniprot database.

### Antibodies

Antibodies used in this study are listed in Table 3.

**Table 3.** List of antibodies used in this study

| Antibody | Company | Applications |
| --- | --- | --- |
| α-taxilin | Proteintech (27558-1-AP) | IF 1:100; IB 1:1000 |
| α-tubulin | Sigma-Aldrich (T9026) | IF 1:1000; IB 1: 10000 |
| β-actin | Cell signaling technology (8H10D10) | IB 1: 1000 |
| γ-taxilin | Proteintech (22357-1-AP) | IF 1:100; IB 1:1000 |
| γ-tubulin | Sigma-Aldrich (T3559) | IF 1:200 |
| Biotin-FITC | Sigma-Aldrich (F6762) | IF 1:100 |
| centrin-3 | Abnova (3E6) (H00001070-M01) | IF 1:200 |
| CEP170 | Abcam (ab72505) | IF 1:100; IB 1:1000 |
|  | Invitrogen (72-413-1) | IF 1:200 |
|  | Proteintech (27325-1-AP) | IF 1:100 |
| EB1 | Proteintech (17717-1-AP) | IF 1:100 |
| FLAG | Sigma-Aldrich (M2) (F3165) | IB 1:10000 |
| GAPDH | ABclonal (AC002) | IB 1:20000 |
| GFP | Laboratory produced rabbit polyclonal antibody | IB 1:1000 |
| GST | Laboratory produced rabbit polyclonal antibody | IB 1: 500 |
| HA | Sigma-Aldrich (H9658) | IF 1:1000; IB 1:1000 |
| ninein | Invitrogen (79.160-7) | IF 1:100; IB 1:1000 |
|  | Abcam (ab231181) | IF 1:100 |
| ODF2 | Proteintech (12058-1-AP) | IF 1:100; IB 1:1000 |
| TCHP | ATLAS antibodies (HPA038638) | IB 1:1000 |
| V5 | Innovative Research (R960CUS) | IF 1:1000; IB 1:10000 |

799 IF, immunofluorescence; IB, immunoblot.

### Gene silencing by siRNA

The siRNAs used in this study (Suzhou GenePharma Co., Ltd) consisted of the following sequences: ODF2, 5’-GGUCACUGUAAAAUGAACC-3’; CCDC68, 5’-CUGCGUGAGUCUUAUUUAU-3’; CCDC120, 5’-GGGAGUGGCUAGUCAUGAU-3’; γ-taxilin #1, 5’-GAAGCAACUGCACAUUUCCAGAUUA-3’; γ-taxilin #2, 5’-GGCAAGAAGCAAGCUAGAAUCUCUU-3’ (*Makiyama et al., 2018*); α-taxilin #1, 5’- GCCUGAACCAACUCCAGUA-3’; α-taxilin #2, 5’-GCGAGGAGCAUAUCGACAA-3’; and α-taxilin #3, 5’-GCGAGGUAUUCACCACAUU-3’. The negative control was set as 5’- UUCUCCGAACGUGUCACGU-3’. All siRNAs were transfected into hTERT RPE1 or HeLa cells by using Lipofectamine 3000 (ThermoFisher Scientific) according to the manufacturer’s instructions, and the cells were analyzed 72 hr post transfection.

### Establishment of α-taxilin and γ-taxilin knockout cell lines

The CRISPR-Cas9 approach was used to generate α- and γ-taxilin KO cell lines (*Ran et al., 2013*). The 20-nt guide sequence of each protein was designed on the online CRISPR Design Tool at http://www.e-crisp.org/E-CRISP/. α-Taxilin was designed as 5’-GCAGTAGAAGCAGAAGGTCC-3’ and γ-taxilin was designed as 5’-CTGTTCGAGCTTCTCCCCCA-3’. The oligo pairs encoding the 20-nt guide sequences were annealed and ligated into a U6-sgRNA plasmid. The U6-sgRNA plasmid, SpCas9-pcDNA3.1, and pcDNA3.1(+)-PuroR were then transfected at a concentration ratio of 2:1:0.5 into HeLa or hTERT RPE1 cells. Puromycin was used to screen or select transfected cells. Clonal cell lines were isolated by Flow cytometry (Astrios EQ) and mutations were detected by sequencing and immunoblotting.

### Immunofluorescence

Cells grown on 18 × 18 mm coverslips were fixed and permeabilized in pre-chilled methanol for 10 min at -20 °C, and then washed three times (10 min each time) in PBS. The cells were then blocked with 4% BSA for 30 min at room temperature, followed by incubation with the first antibody (diluted in 4% BSA) at 4 °C overnight. The antibodies used for immunofluorescence are listed in Table 1. The slides were then washed three times in PBS buffer for 10 min each time, blocked with 4% BSA for 30 min at room temperature, and then incubated with the second antibody (Alexa Fluor, Invitrogen) for 1 hr at room temperature. Finally, the cells were stained with DAPI.

Confocal and STED images were acquired using a TCS SP8 STED 3X (Leica) with a 100× 1.4 NA APO oil objective lens and LAS X software 2.0 (Leica), or STEDYCON (Abberior Instruments). Huygens software 14.10 (Scientific Volume Imaging, the Netherlands) was used for STED nanoscopy image deconvolution. An N-SIM Microscope system equipped with a 100× 1.49 NA APO oil objective lens (Nikon) was used to acquire 3D-SIM images. NIS-Elements AR software 4.51 (Nikon) was used both to acquire the images and for three-dimensional reconstruction. Images were then processed in Photoshop (Adobe).

### Immunoprecipitation and immunoblots

For immunoprecipitation, HEK-293T or hTRET RPE1 cells were lysed in immunoprecipitation lysis buffer (50 mM HEPES, 250 mM NaCl, 0.1% Nonidet P-40, 1 mM DTT, 1 mM EDTA, and 10% glycerol, pH = 7.4) on ice for 30 min and then the lysates were centrifuged at 12, 000*g* for 15 min at 4 °C. Then, Protein A-Sepharose beads or Protein G Sepharose beads (Amersham Biosciences) (Amersham Biosciences) were added to the supernatants, which were then individually incubated with appropriate antibodies (*Table 1*) overnight at 4 °C. The beads were then washed with immunoprecipitation lysis buffer and collected in SDS loading buffer (50 mM Tris-HCl, pH = 7.4, 2% SDS, 100 mM DTT, 0.025% bromo blue, 10% glycerol), which was then boiled at 100 °C for 10 min to get the immunoprecipitation samples.

For immunoblots, SDS-PAGE was used to separate protein samples, which were then transferred onto polyvinylidene difluoride membranes (#IPVH00010, Millipore). The membranes were blocked with 5% non-fat milk for 30 min, and then incubated with primary antibodies (listed in Table 1) overnight at 4 °C, or for 2 h at room temperature. After incubation with primary antibodies, the membranes were then incubated with peroxidase-affinipure goat anti-rabbit or mouse IgG (H+L) secondary antibodies (#111-035-003 and #115-035-003, Jackson ImmunoResearch, 1:5000) for 1 hr at room temperature.

### GST pull down assay

For GST pull-down assay, bacterial-expressed GST and GST-α-Taxilin-M (expessed in *Escherichia coli* strain BL21) were bound to glutathione-Sepharose 4B beads (GE Healthcare) and incubated with bacterial-expressed MBP-γ-Taxilin-M or 293T cell expressed and purified 3×FLAG-CEP170 at 4 °C overnight. After reaction, complexes were washed at least 5 times with GST-binding buffer (20 mM HEPES, pH 7.5, 75 mM KCl, 0.1 mM EDTA, 2.5 mM MgCl2, 1 mM DTT, 0.05% Triton X-100), eluted by boiling in SDS-PAGE loading buffer, and subjected to immunoblot with the indicated antibodies.

### Microtubule regrowth assay

RPE-1 cells grown on 18 × 18 mm coverslips and then the coverslips were embedded on ice for 30 min to depolymerize the cytoplasmic microtubules. The cells were then brought back to 37 °C to allow the microtubules to reform. Cells at 0, 5, and 10 min after rewarming were fixed in 4% PFA (pre-warmed to 37 °C) for 10 min at 37 °C. The cells were immunostained with α- and γ-tubulin antibodies.

### Measurements and statistical analysis

Immunofluorescence intensities and the diameters of the ring-like SDA protein structures were measured using Image J software 1.48v (NIH). The statistical significance among different groups was determined by two-tailed student’s *t*-tests or one-way ANOVA. The data were graphed in Prism 5 (GraphPad).

## Acknowledgments

We thank the Flow Cytometry Core at the National Center for Protein Sciences at Peking University in Beijing, China, particularly Liying Du, Jia Luo, Huan Yang, and Hongxia Lv, for their technical help. We also thank the Optical Imaging Core Facility at the National Center for Protein Sciences, particularly Chunyan Shan and Ye Liang for assistance with the 3D-SIM, confocal, and STED microscopy imaging. We thank all members of the laboratory for their helpful discussions. This work was supported by the National Natural Science Foundation of China (31630092, 31830110, and 31801133) and the National Key Research and Development Program of China, Stem Cell and Translational Research (2016YFA0100501).

## Authors contributions

Dandan Ma planned the project, performed the experiments, and wrote the manuscript. Rongyi Wang performed the molecular cloning, Fulin Wang processed all the images, Zhiquan Chen and Yuqing Xia performed statistical analysis. Ning Huang modified the manuscript. Junlin Teng and Jianguo Chen supervised the work.

---

**Figure 1–figure supplement 1.**
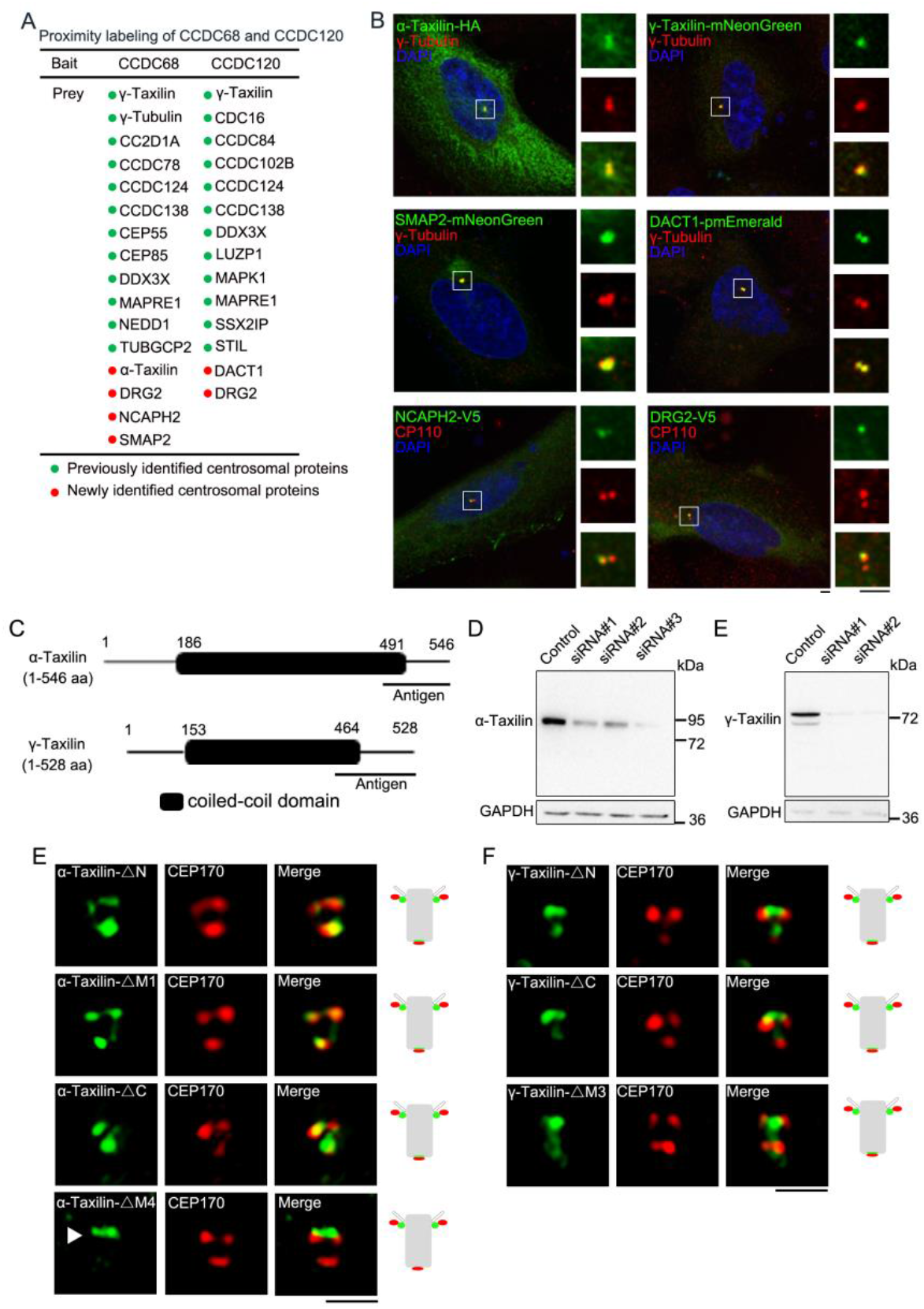
Screen of α-Taxilin and γ-taxilin as subdistal appendage proteins by CCDC68 and CCDC120 proximal labeling, and their localization characteristics. **(A)** Centrosomal proteins found using mass spectrometry examination of APEX2-mediated biotin proximal labeled CCDC68-V5-APEX2 and CCDC120-V5-APEX2. **(B)** Confocal images of the indicated infusion proteins (green) co-localized with immunostained centrosome markers CP110 (red) or γ-tubulin (red) (arrows) in U2OS cells. DNA was stained with DAPI (blue). Scale bars, 2.5 µm. **(C)** α-Taxilin and γ-Taxilin schematic showing their N-terminus, M region (coiled-coil domain), and C-terminus. **(D)** Immunoblots showing the anti-α-taxilin antibody specificity in the control and α-taxilin siRNA-treated U2OS cells. GAPDH was the loading control. **(E)** Structured illumination microscopy (SIM) images of immunostained HA-tagged α-taxilin deletion mutants (green) and CEP170 (red) in RPE-1 cells. Scale bar, 1 µm. **(F)** SIM images of immunostained CEP170 (red) in RPE-1 cells transfected with GFP-tagged γ-taxilin deletion mutants (green). Scale bar, 1 µm. The cartoons to the right of the images in (E) and (F) graphically depict the merge images.

**Figure 3–figure supplement 1.**
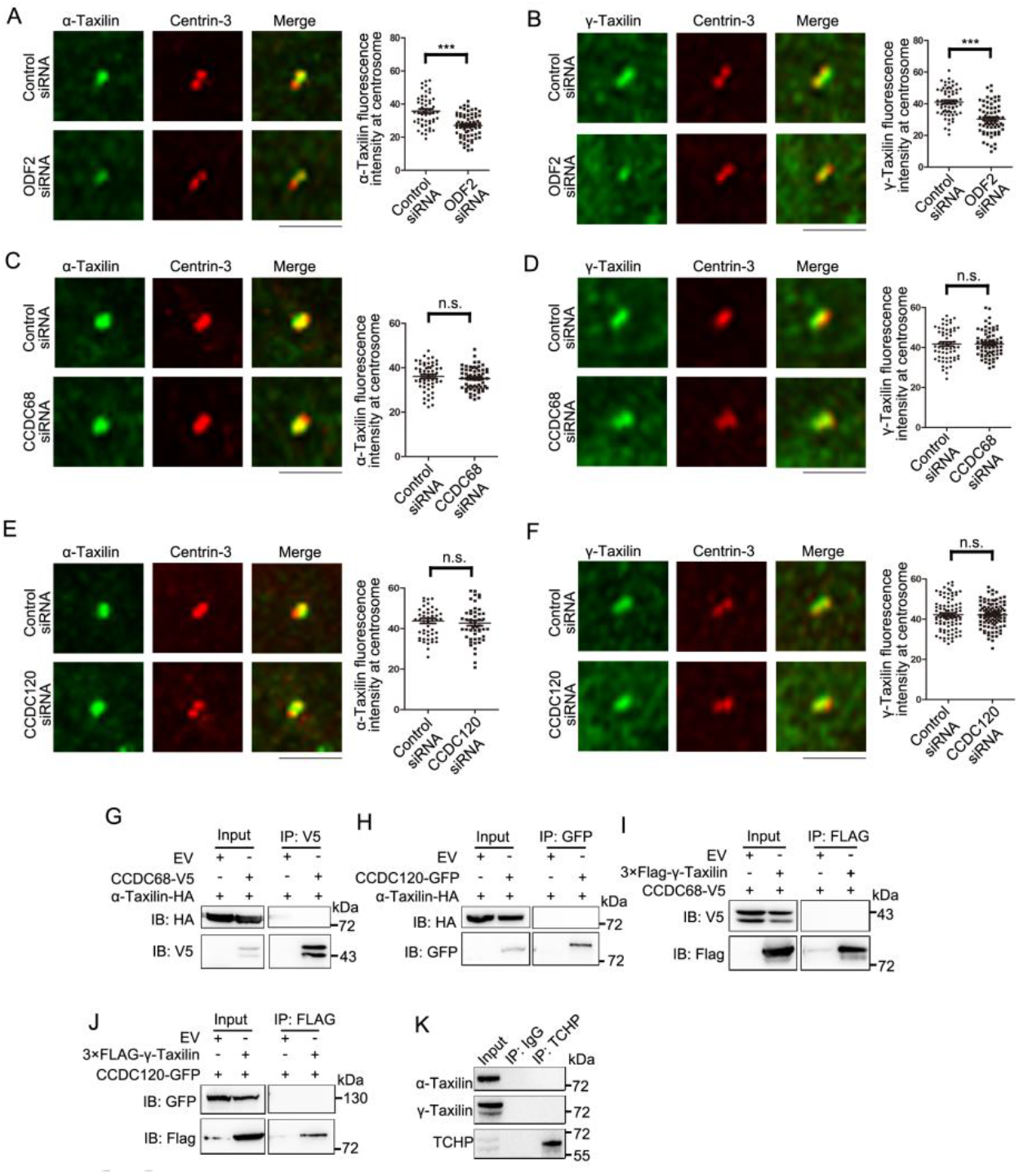
α-Taxilin and γ-taxilin assembly at the centrosome. **(A-B)** Confocal images and statistical analyses of immunostained α-taxilin (A) and γ-taxilin (B) (green) at the centrosome in control- or ODF2-siRNA treated RPE-1 cells. Immunostained centrin-3 (red) served as the centrosome marker. **(C-D)** Confocal images and statistical analysis of α-Taxilin (C) and γ-Taxilin (D) (green) at the centrosome in control- or CCDC68-siRNA treated RPE-1 cells. Immunostained centrin-3 (red) served as the centrosome marker. **(E-F)** Confocal images and statistical analyses of α-taxilin (E) and γ-taxilin (F) (green) at the centrosome in control- or CCDC120-siRNA treated RPE-1 cells. Immunostained centrin-3 (red) served as the centrosome marker. For (A-F), scale bars, 5 µm. Data are mean (SEM). Statistical significance was determined by the two-tailed Student’s *t*-tests. ***, *p* < 0.001; n.s., not significant. **(G)** Lysates of HEK-293T cells co-overexpressing CCDC68-V5 and α-taxilin-HA were subjected to immunoprecipitation (IP) with anti-V5 antibody and immunoblotted (IB) with anti-V5 and anti-HA antibodies. **(H)** Lysates of HEK-293T cells co-overexpressing CCDC120-GFP and α-taxilin-HA were subjected to IP with anti-GFP antibody and IB with anti-GFP and anti-FLAG antibodies. **(I)** Lysates of HEK-293T cells co-overexpressing CCDC68-V5 and 3×FLAG-γ-taxilin were subjected to IP with anti-FLAG and IB with anti-V5 and anti-FLAG antibodies. EV, empty vector. **(J)** Lysates of HEK-293T cells co-overexpressing CCDC120-GFP and 3×FLAG-γ-taxilin were subjected to IP with anti-FLAG and IB with anti-GFP and anti-FLAG antibodies. **(K)** Endogenous IP assay of α-taxilin or γ-taxilin with TCHP using anti-TCHP antibody in lysates of HEK-293T cells and IB with anti-α-taxilin, anti-γ-taxilin, and anti-TCHP antibodies. IgG was the control.

**Figure 4–figure supplement 1.**
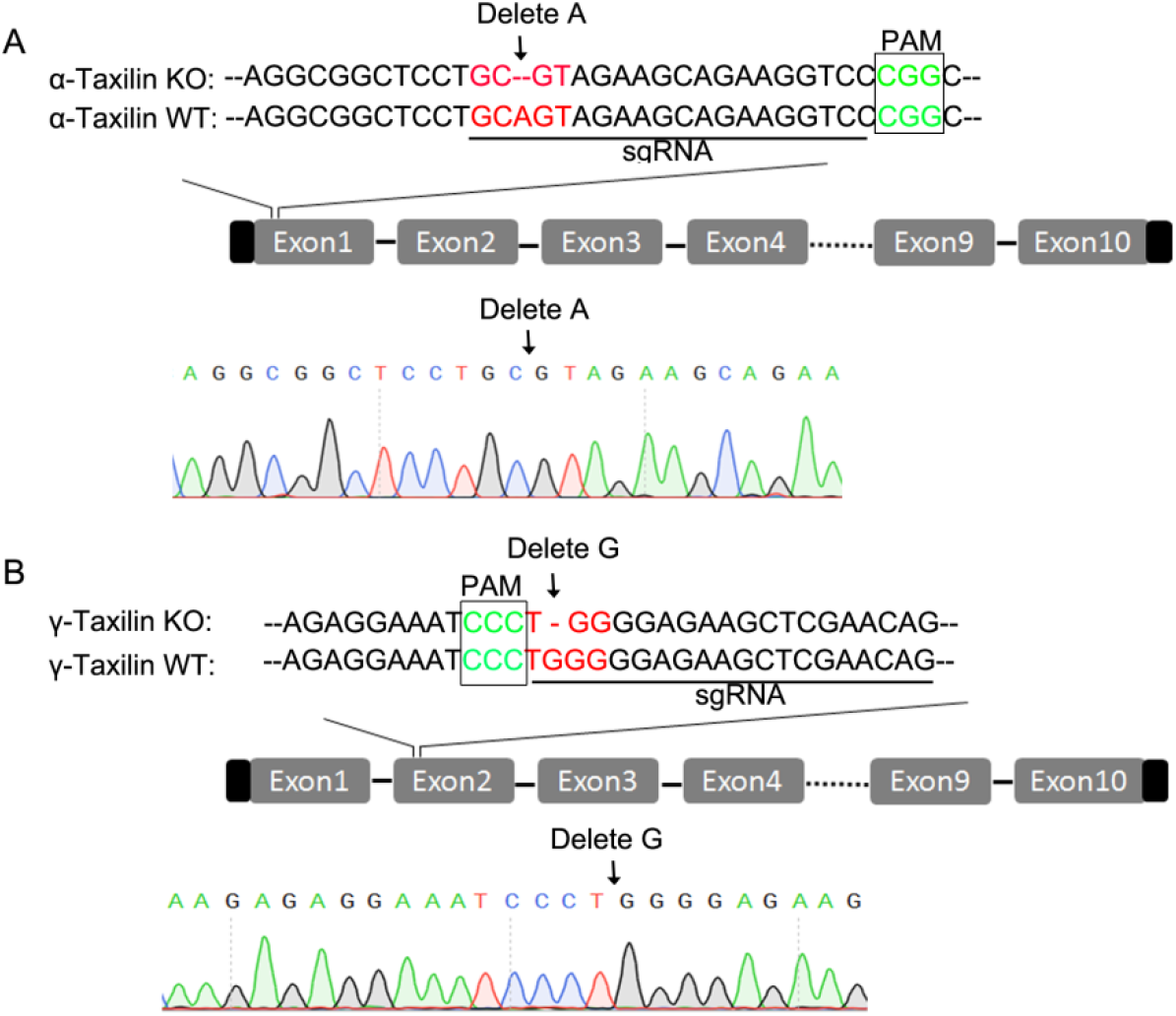
Partial sequences of *α-taxilin* or *γ-taxilin* knockout (KO) in RPE1 cells. **(A-B)** Sequence alignments of partial *α-Taxilin* (A) and *γ-Taxilin* (B) coding sequences from their wild-type (WT) and KO RPE-1 cells. The protospacer adjacent motif (PAM) sequences are in green font, the sgRNA target sequences are underlined, and mutated sequences are in red font.

**Figure 6–figure supplement 1.**
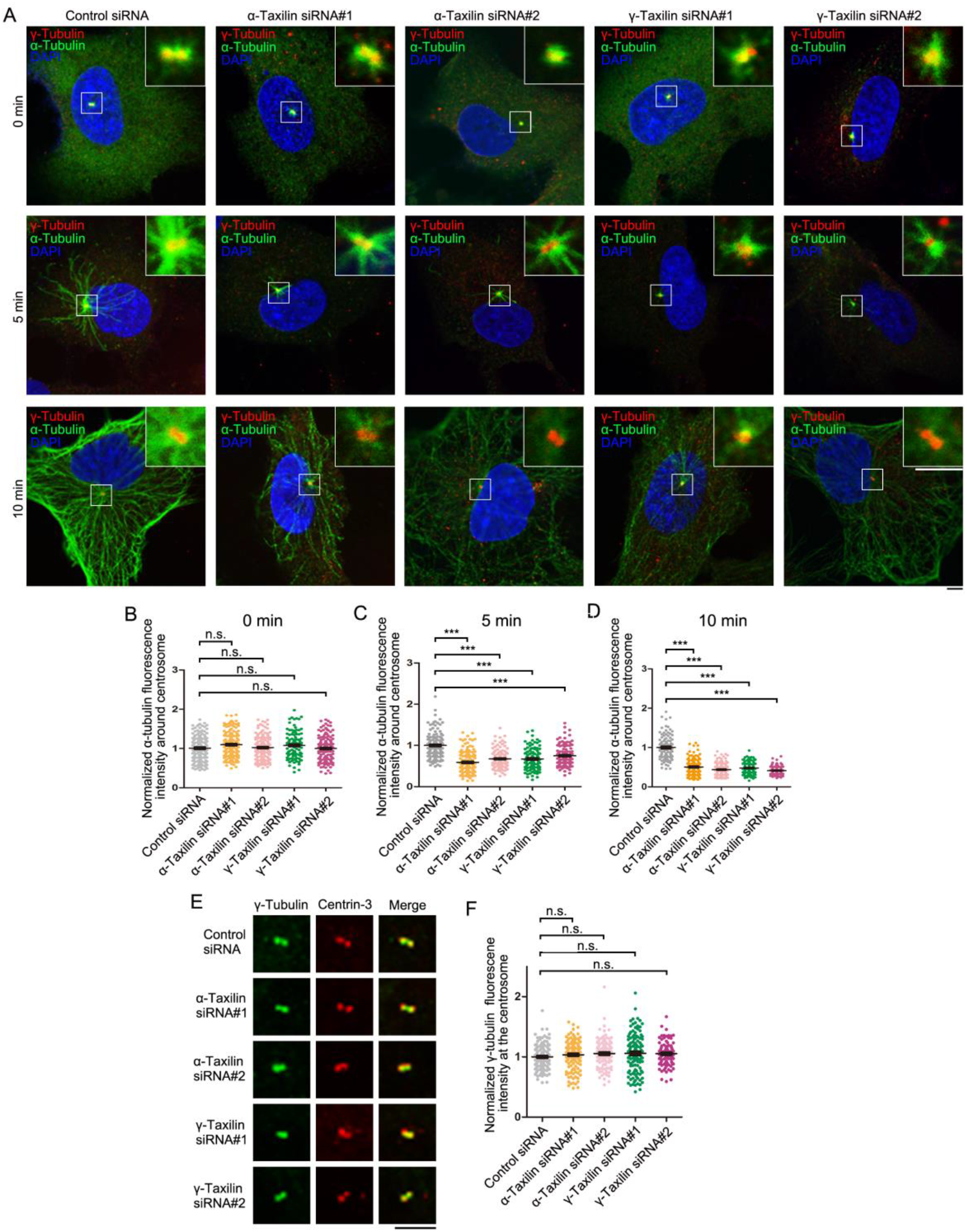
α-Taxilin and γ-taxilin depletion inhibits microtubule reformation in RPE-1 cells after cold depolymerization. **(A)** Confocal images of microtubule reformation with immunostained α-tubulin (green) in control-, α-Taxilin- or γ-Taxilin-siRNA treated RPE-1 cells. Immunostained γ-tubulin (red) was used as the centrosome marker. DNA was stained with DAPI (blue). Scale bars, 10 µm. **(B-D)** Statistical significance of α-tubulin fluorescence intensity at 0 min (B), 5 min (C), and 10 min (D) was determined by one-way ANOVA of three independent experiments with over 100 cells total. Data are mean (SEM). ***, *p* < 0.001; n.s., not significant. **(E)** Confocal images of immunostained γ-tubulin (green) and centrin-3 (red) in control-, α-taxilin- or γ-taxilin-siRNA treated RPE-1 cells. Scale bar, 5 μm. **(F)** Comparisons of γ-tubulin fluorescence intensity at the centrosomes in (E). Statistical significance was determined by one-way ANOVA. Data are mean (SEM). n.s., not significant.

**Figure 7–figure supplement 1.**
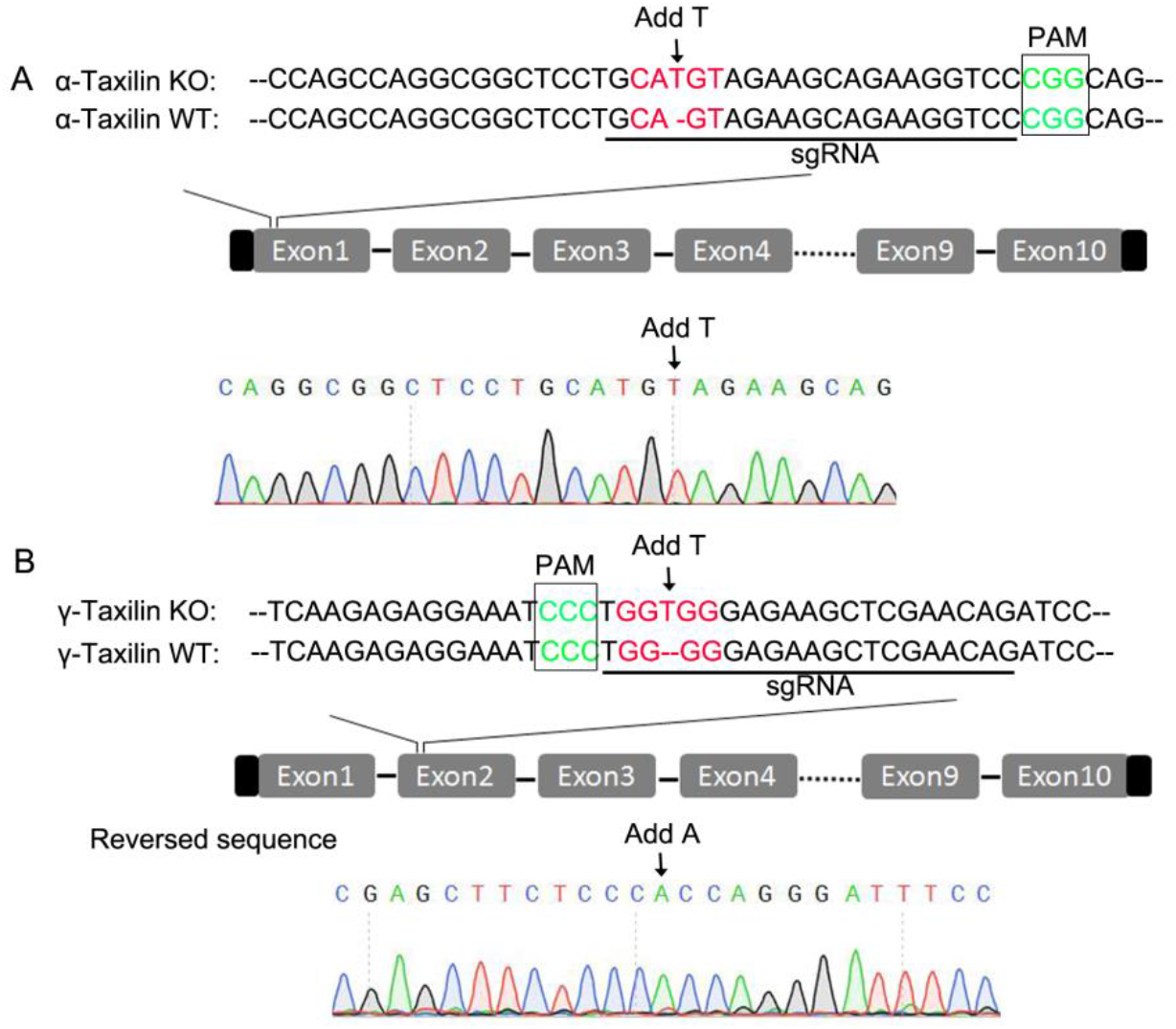
Partial sequences of *α-taxilin* or *γ-taxilin* knockout (KO) in HeLa cells. **(A-B)** Sequence alignments of partial *α-Taxilin* (A) and *γ-Taxilin* (B) coding sequences from their wild-type (WT) and KO HeLa cells. The protospacer adjacent motif (PAM) sequences are in green font, the sgRNA target sequences are underlined, and mutated sequences are in red font.

